# Large-scale genomic rearrangements boost SCRaMbLE in *Saccharomyces cerevisiae*

**DOI:** 10.1101/2023.05.21.541650

**Authors:** Tianyi Li, Shijun Zhao, Li Cheng, Sha Hou, Zhouqing Luo, Jinsheng Xu, Wenfei Yu, Shuangying Jiang, Marco Monti, Daniel Schindler, Weimin Zhang, Chunhui Hou, Yingxin Ma, Yizhi Cai, Jef D. Boeke, Junbiao Dai

## Abstract

Genomic rearrangements contribute to gene copy number alterations, disruption of protein-coding sequences and/or perturbation of cis-regulatory networks. SCRaMbLE, a Cre/loxP-based system implanted in synthetic yeast chromosomes, can effectively introduce genomic rearrangements, and is thus a potential tool to study genomic rearrangements. However, the potential of SCRaMbLE to study genomic rearrangements is currently hindered, because a strain containing all 16 synthetic chromosomes is not yet available. Here, we constructed a yeast strain, SparLox83, containing 83 loxPsym sites distributed across all 16 chromosomes, with at least two sites per chromosome. Inducing Cre recombinase expression in SparLox83 produced versatile genome-wide genomic rearrangements, including inter-chromosomal events. Moreover, SCRaMbLE of the hetero-diploid strains derived from crossing SparLox83 with strains possessing synthetic chromosome III (synIII) from the Sc2.0 project led to increased diversity of genomic rearrangements and relatively faster evolution of traits compared to a strain with only synIII. Analysis of these evolved strains demonstrates that genomic rearrangements can perturb the transcriptome and 3D genome structure and can consequently impact phenotypes. In summary, a genome with sparsely distributed loxPsym sites can serve as a powerful tool to study the consequence of genomic rearrangements and help accelerate strain engineering in Saccharomyces cerevisiae.

## Introduction

Genomic rearrangements alter genetic linkage of discrete chromosomal regions and are an important mutational force in evolution^1, 2^, underlying the pathology of various germline and somatic diseases such as nervous system disorders^3^, congenital heart diseases^4^, and cancers^5^. The profound effects of these genomic rearrangements, which include deletions, duplications, insertions, inversions, and translocations, are attributed to alterations in gene copy numbers, disruption of protein-coding sequences, and perturbation of cis-regulatory networks^6–10^. Previous studies indicate that the rate of genomic rearrangements is several orders of magnitude higher than the rate of base substitutions^11^, and genomic rearrangements are pervasive in all domains of life, from prokaryotes to humans^12^.

Genomic rearrangements likely occur randomly and sporadically, and discriminating their impacts from the confounding effects of other mutation types remains a challenge^13, 14^. To overcome this problem, several methods have been developed to construct targeted long-range rearrangements in model organisms, including *I-Sce*I-induced DSB repair^15^, Cre/loxP recombination^13^, and CRISPR/Cas9^14^. However, only limited and pre-determined inter-chromosomal rearrangements can be generated and analyzed by these methods. A controlled system allowing genome-wide induction of a wide array of chromosomal rearrangements has not yet been established.

In the synthetic yeast genome project (Sc2.0), thousands of loxPsym sequences are incorporated in a designer genome to create a system known as <u>S</u>ynthetic <u>C</u>hromosome <u>R</u>ecombination <u>a</u>nd <u>M</u>odification <u>b</u>y <u>L</u>oxP-mediated <u>E</u>volution (SCRaMbLE)^16^. Through SCRaMbLE, massive rearrangements can be introduced into the synthetic chromosomes, including intra-chromosomal deletions, inversions, duplications, and inter-chromosomal translocations^17, 18^. SCRaMbLE therefore provides a strategy to investigate the potential mechanisms of genomic rearrangements and their functional consequences and facilitates delineation of the relationships between large-scale genomic rearrangements and diseases. However, to date, the power of SCRaMbLE has mostly been used in single synthetic chromosome contexts^18–30^. Moreover, intra-chromosomal rearrangements dominate, with inter-chromosomal rearrangements rarely detected in individual SCRaMbLEd strains^31^.

Here, CRISPR/Cas9 was used to simultaneously integrate loxPsym sequences into multiple genomic loci. The resultant extensively engineered yeast strain, SparLox83, possessed <u>83</u> <u>Spar</u>sely distributed <u>L</u>oxPsym sites across the genome. After inducing expression of Cre recombinase, the resultant SCRaMbLEd yeast population underwent diverse large-scale genomic rearrangements, dominated by inter-chromosomal events. By quantifying rearrangement frequencies at loxPsym sites, this study revealed multiple factors impacting loxPsym-mediated genomic rearrangements, including chromatin accessibility and genomic distance between sites. The impacts of large-scale genomic rearrangements on cell fitness under stress were investigated in progeny populations, and translocation and duplication events were detected that led to increased tolerance of nocodazole. SparLox83 was further enhanced with the SCRaMbLE system by crossing with a haploid strain containing synIII (JDY541) from the Sc2.0 project. Inducing SCRaMbLE in this heterozygous diploid strain resulted in complex genomic rearrangements, including numerous loss of heterozygosity (LOH) and aneuploidy events, as well as rapid adaptation of tolerance to acetic acid (HAc). The loxPsym sites in SparLox83 support the rapid development of new phenotypes by substantially boosting the SCRaMbLE system with large-scale genomic rearrangements.

## Results

### Genome-wide insertion of 83 loxPsym sites throughout the yeast genome

Cre/LoxP based SCRaMbLE cannot yet be employed in the Sc2.0 project across all 16 yeast chromosomes because construction of a fully synthetic yeast genome is still underway. Previous studies employing SCRaMbLE in strains with synthetic chromosomes reported the frequency of inter-chromosomal rearrangements as less than 10%^31, 32^. To construct an engineered yeast strain suitable for effective introduction of genome-wide, large-scale genomic rearrangements, loxPsym-containing sequences were introduced by design into all the yeast chromosomes using a CRISPR-Cas9 based gene editing strategy (Extended Data Fig. 1a). Briefly, three loci were simultaneously targeted for integration of loxPsym sites in each cycle, eventually producing a strain with 83 sites (JDY525) distributed across the 16 chromosomes, with at least two sites per chromosome (Fig. 1a). The presence of loxPsym at each locus was confirmed by whole genome sequencing (WGS; Table S1). LoxPsym sites in JDY525 were sparsely distributed, with significantly longer genomic distances between adjacent sites than in Sc2.0 synthetic chromosomes. The average genomic distance between adjacent LoxPsym sites was approximately 145 kb in JDY525, much larger than the 3 kb distance between sites in Sc2.0 chromosomes (Extended Data Fig. 1b). Furthermore, WGS and pulsed-field gel electrophoresis (PFGE) revealed a chromosome translocation between chrVIII and chrIX in JDY525 (Extended Data Fig. 2).

**Fig. 1.**
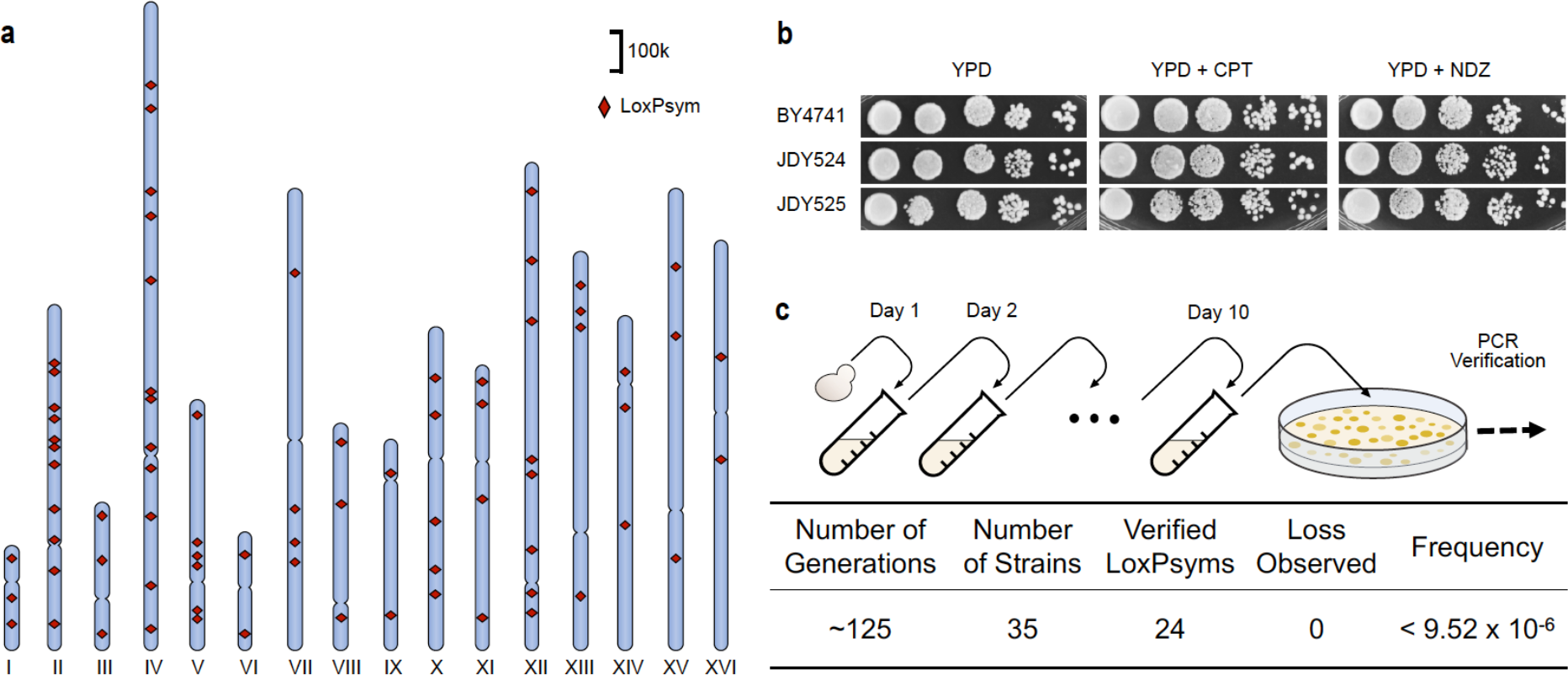
Optimization, construction, and characterization of a yeast strain with loxPsym sites distributed throughout the genome. **a**. Distribution of 83 loxPsym sites in the JDY525 strain. Detailed site locations are in Table S1. **b**. JDY525 phenotyping on various media. Ten-fold serial dilutions of overnight cultures of JDY525, parental strain JDY524, and wild-type BY4741 were analyzed on YPD, YPD + camptothecin (CPT, 15 μg/mL), and YPD + nocodazole (NDZ, 2.5 μg/mL). YPD, yeast extract peptone dextrose. Images taken after two days incubation at 30°C. **c.** PCR analysis of JDY525 after approximately 125 generations to assess genomic stability. Potential loss of loxPsym was evaluated at 24 different loci in 35 passaged strains and no losses were observed. The frequency is a maximum estimate of loxPsym loss frequency per generation.

**Fig. 2.**
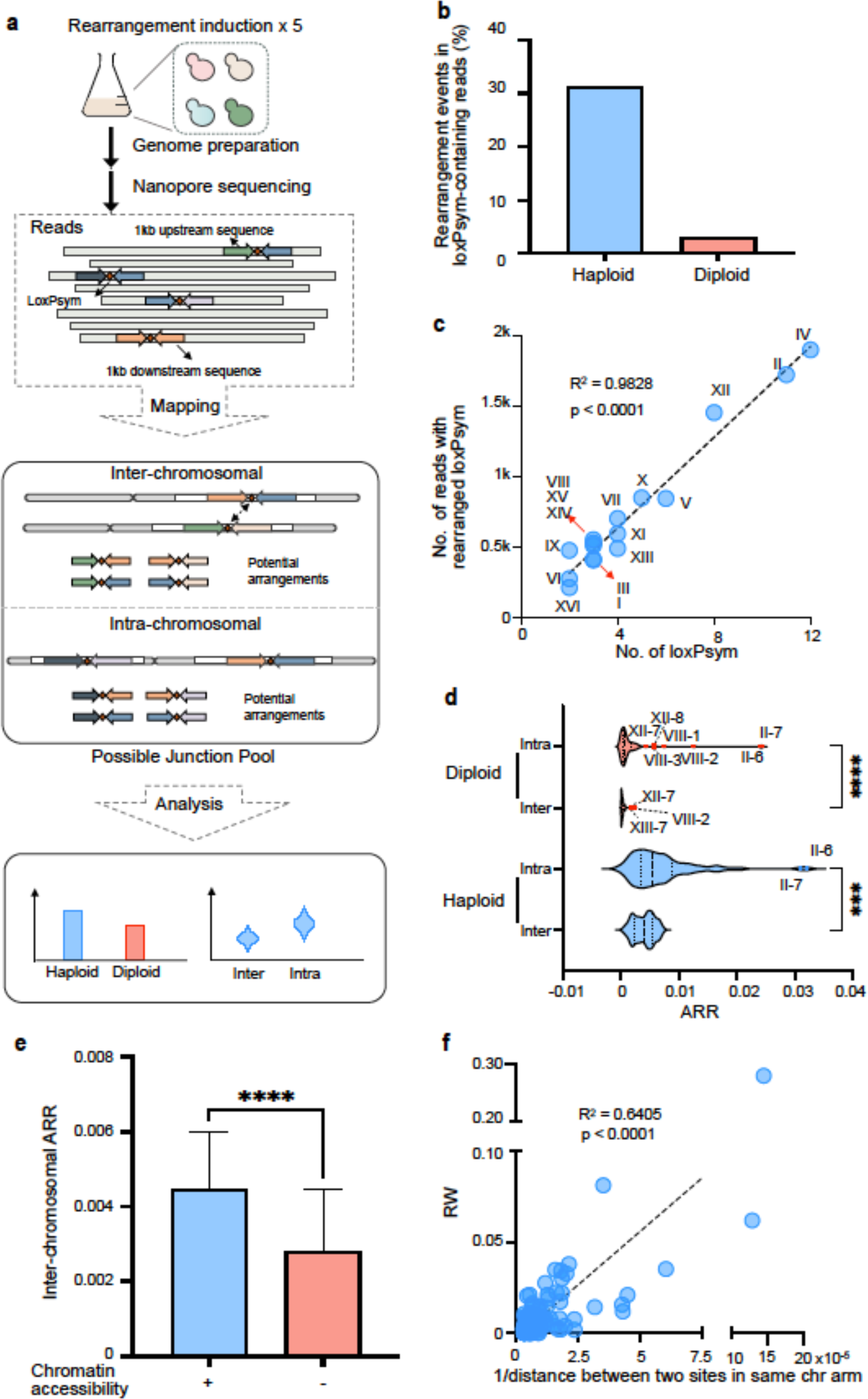
Characterization of genome-wide rearrangement by SCRaMbLE. **a**. Workflow for loxPsym-mediated genome-wide rearrangement and analysis. A single colony was cultured and subjected to five sequential rounds of Cre recombinase induction to generate a SCRaMbLEant pool. The genomic DNA was then sequenced. Potential rearrangements were identified and analyzed by mapping the loxPsym-containing reads to the sequence pool comprising all possible pairwise combination sequences 1 kb upstream (forward arrow) and 1 kb downstream (backward arrow) of each loxPsym site. **b**. Percentage of reads exhibiting genomic rearrangements in detected loxPsym-containing reads in haploid and diploid SCRaMbLEant cells. **C**. Correlation between number of identified rearrangements and number of loxPsym sites in each chromosome. Y-axis, number of reads with rearrangement; X-axis, number of loxPsym sites in a chromosome. **d**. Average rearrangement rate (ARR) in haploid and diploid cells. *p < 0.05; **p < 0.01; ***p < 0.001; ****p < 0.0001. **e**. Comparison of inter-chromosomal ARRs between loxPsym sites within (+) and outside (-) open chromatin regions in haploid cells; ****p < 0.0001. **f**. Correlation between Rearrangement Weight (RW) and 1/genomic distance of loxPsym sites in the same chromosome arm in haploid cells. R^2^=0.6405 and p<0.0001.

The fitness of JDY525 was evaluated by comparing growth in various conditions with its parental strain JDY524 (without loxPsym) and BY4741 in serial dilution tests. Growth of JDY525 was indistinguishable from JDY524 or BY4741 on YPD and under stress conditions, including the presence of 15 μg/ml camptothecin (an inhibitor of topoisomerase I that causes genome instability) or 2.5 μg/ml nocodazole (a drug that affects mitosis by interference with microtubule polymerization) (Fig. 1b). Given the presence of multiple loxPsym sites in its genome, the genome stability of JDY525 was checked using a previously described junction PCR-based assay^33^. None of the loxPsym sites were lost in 35 independent lineages after ∼120 mitotic generations on YPD in the absence of Cre (Fig. 1c). These results confirmed the successful insertion of 83 loxPsym sites into the wild-type yeast genome without impairment of genome stability or fitness under various conditions.

### Induction of Cre recombinase activity in SparLox83 leads to versatile rearrangements, particularly inter-chromosomal rearrangements

To facilitate the selection of clones with genomic rearrangements, the previously described ReSCuES system^26^ was integrated into JDY525 to generate SparLox83. Since there is at least one essential gene positioned between each pair of adjacent loxPsym sites, deletions will result in cell death, limiting the extent of all the possible rearrangements. Therefore, an additional heterozygous diploid strain, JDY536, was generated by crossing SparLox83 with BY4742^34^.

SparLox83 and JDY536 were induced to express Cre recombinase and selected using the ReSCuES system for five consecutive rounds before collection of SCRaMbLEants. No other selection was imposed. First, to confirm the presence of rearrangements, PCR was used to amplify some of the novel junctions that were not present in the original SparLox83 genome but which might be generated following loxPsym-derived rearrangements (Extended Data Fig. 3a). Next, the pooled SCRaMbLEants were subjected to Nanopore MinION sequencing. To avoid PCR bias and obtain reads that were as long as possible, a ligation-sequencing strategy with no PCR amplification was used, and genome shredding steps were avoided. In total, 774,791 (8.06 Gb) and 2,523,754 (22.5 Gb) reads were generated from the SparLox83 and SparLox83 × BY4742 SCRaMbLEant populations, respectively.

A population-based strategy was adopted to detect and characterize rearrangement events in SCRaMbLEant populations (Fig. 2a). Flanking regions of loxPsym sites were extracted and assembled with all potential rearrangements *in silico*. Next, Nanopore reads containing loxPsym sequences were aligned to the potential rearrangements. Each filtered match between a read and a potential rearrangement was defined as a rearrangement event. This analysis revealed that 5721 and 936 rearrangement events took place across all loxPsym sites in the haploid and diploid strains, which accounted for 31.1% and 3.1% detected loxPsym-containing reads, respectively (Fig. 2b). Flanking region lengths were set to 500, 1000, 1500, and 2500 bp, with no significant differences seen in the analysis results (Extended Data Fig. 3b). The following results are those obtained using 1000 bp flank regions. At least one rearrangement event was detected for every loxPsym site in both the haploid and diploid strains. Inter-chromosomal rearrangements predominated: 4852 (84.8%) and 595 (63.6%) of the rearrangement events were inter-chromosomal in haploid and diploid cells, respectively. Rearrangements were next assigned to their different chromosomes to test whether rearrangement events occurred evenly across the genome. In haploid cells, an increased number of rearrangement events was associated with a higher number of loxPsym sites on that chromosome (Fig. 2c, r=0.9828, p<0.001).

These results abtaoned in the absence of selection beyond the requirement for at least one RESCUES inversion event per cell showed that genome-wide insertion of loxPsym sites in SparLox83 produced versatile rearrangements, with a bias towards inter-chromosomal events, upon Cre induction. The preference for inter-chromosomal rearrangements suggests that SparLox83 presents an ideal model for studying the impacts of large-scale genomic rearrangements.

### Quantitative analysis revealed an uneven distribution of rearrangements at different loxP sites

Previous studies used rearrangement event frequencies at particular loxPsym sites to quantify their rearrangement capacities^31, 32^. Rearrangement event frequencies were, however, biased by the different sequencing depths at each loxPsym site. To overcome this, an Average Rearrangement Rate (ARR), a normalized rearrangement event frequency, was used to quantify the average rearrangement potential for each loxPsym site. ARRs were compared for intra- and inter-chromsomal rearrangements. Although intra-chromosomal ARRs were significantly higher than inter-chromosomal ARRs, the difference between the two categories was limited (Fig. 2d). Moreover, outlier ARRs represented particularly active sites. Seven outliers (II-7, II-6, VIII-2, VIII-1, XII-8, VIII-3, XII-7) were found for intra-chromosomal ARRs in diploid cells, two of which (II-7 and II-6) were also observed in haploid cells. Three sites (VIII-2, XII-7, XIII-7) were identified as inter-chromosomal outliers in diploid cells, but no outliers were observed for inter-chromosomal rearrangements in haploid cells.

Rearrangement Weight (RW) was derived to quantitatively describe the rearrangement event frequency between two specific sites and reduce the bias caused by sequencing depth. RW divides the number of rearranged events by the geometric mean of the sequencing depths of pairs of interacting sites. The RW matrix containing RWs of all possible loxPsym site pairs (2775 pairs) was generated. Of these, 2111 (76.1%) non-zero RWs were represented in haploid cells, suggesting that most of the potential rearrangement events occurred. Interaction maps of the RW matrix for intra- and inter-chromosomal rearrangements were generated (Extended Data Fig. 4). The RW between sites II-6 and II-7 (0.28) was significantly higher than other RW values, which were below 0.087 (Extended Data Fig. 4a). Inter-chromosomal RWs also exhibited an uneven distribution (Extended Data Fig. 4b), with events between site pairs located in the inner circle (RWs>0.02) occurring more frequently than with other site pairs (RWs<0.02). Together, these results showed that loxPsym sites in SparLox83 exhibited highly variable rearrangement event frequencies.

### The genomic locations of loxPsym sites impact their rearrangement event frequencies

Previous studies using SCRaMbLE in strains harboring one or more synthetic chromosomes confirmed that rearrangement frequency varied among loxPsym sites^31, 32^. In synthetic chromosomes, where loxPsym sites are densely arranged, the rearrangement frequencies of loxPsym sites correlated with chromatin accessibility and spatial contact probability^32^. However, due to the preference for intra-chromosomal rearrangements in these synthetic chromosomes, factors impacting inter-chromosomal rearrangement frequencies remain less well understood. The genome-wide distribution of loxPsym sites in SparLox83, with their frequent inter-chromosomal rearrangements and quantitatively defined RWs, indicate that SpaL83 is an ideal model to explore the features of rearrangement frequencies involved in sparsely arranged loxPsym sites.

Consistent with previous findings^32^, the inter-chromosomal ARRs of sites positioned at open chromatin regions^35^ were significantly higher than sites outside open chromatin regions (Fig. 2e).

The highest RW was observed between sites II-6 and II-7, which possessed the shortest genomic distance among the 83 loxPsym sites. Next, the impact of genomic distance between two intra-chromosomal loxPsym sites on RW was examined. The RW between two loxPsym sites located in the same chromosome arm showed a negative correlation with genomic distance in haploid and diploid cells (Fig. 2f, Extended Data Fig. 5a). For two sites not on the same chromosome, spatial distance was determined using genome-wide chromosome conformation capture analysis (Hi-C), which facilitated the generation of a normalized contact map of the SparLox83 genome (Extended Data Fig. 5b). However, no correlation was found between the normalized contact counts and the RW of paired loxPsym sites.

Together, these results suggest that chromatin accessibility and genomic distance partly impact the rearrangement frequency of loxPsym sites.

### Large-scale genomic rearrangement and stress tolerance

Large-scale rearrangements drive phenotypic diversification and environmental adaptability^36, 37^, but distinguishing the impact of large-scale rearrangements on fitness from other types of mutation in naturally evolved strains remains challenging. The SparLox83 strain can be used to introduce large-scale genomic rearrangements, allowing the investigation of the role of such rearrangements in cell fitness alterations under various conditions.

Rearrangements were induced in SparLox83 and strains were screened under a range of selective stresses including high alcohol (10% ethanol), high oxidation (3 mM H2O2), high osmotic stress (1 M NaCl), and non-fermentable sugar as carbon source (3% glycerol). Experiments were repeated in the presence of reagents to perturb microtubules (10 μg/ml nocodazole and 40 μg/ml benomyl), cause DNA damage (30 μg/ml camptothecin), or inhibit the TOR pathway (10 ng/ml rapamycin) (Fig. 3a) (Luo et al., 2018). Although no candidates conferring increased tolerance to ethanol, H_2_O_2_, or NaCl were identified, several clones were more resilient to nocodazole, benomyl, or rapamycin than the original strain (Fig. 3b and Extended Data Fig. 6a).

**Fig. 3.**
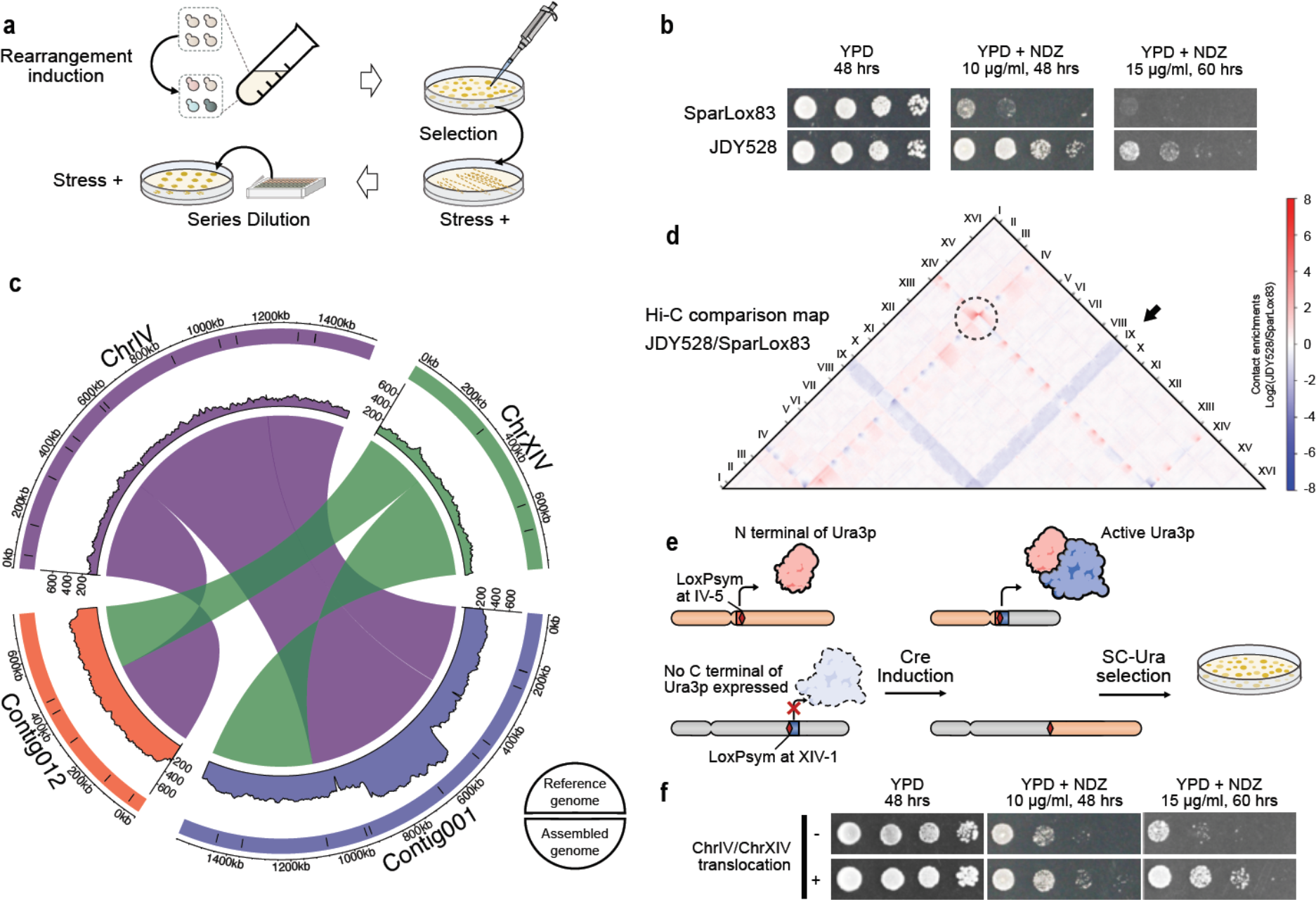
Genome-wide rearrangements confer resistance to various stresses. **a.** Workflow used to identify strains tolerant to various stresses. A single colony was cultured and subjected to rearrangement induction before plating onto media containing drug, alcohol, oxidation, or osmotic stress treatments. Colony phenotypes were verified using serial dilution assays. **b**. Ten-fold serial dilutions of overnight cultures of the original strain (SparLox83) and a nocodazole-resistant clone (JDY528) analyzed on YPD and YPD containing different concentrations of nocodazole (YPD+NDZ). **c**. Colinearity mapping between assembled contig001 (blue) and contig012 (red) in JDY528 with chrIV (purple) and chrXIV (green) of the reference genome. The outer circle represents chromosomes before (upper half) and after (lower half) translocation. Black bars in the outer circle indicate the locations of loxPsym sites. The middle circle is a depth plot of the Nanopore sequencing data, and the inner circle indicates the inferred structural variants. **d**. Ratio plots of contact maps of the SparLox83 and JDY528 genomes showing translocations (dotted circle) and chromosome position changes (black arrow). **e**. Representation of the split-*URA3* method used to reconstruct the chrIV/chrXIV translocation in JDY524 (*i.e.*, the parental strain lacking loxPsym sites). **f**. Ten-fold serial dilutions of strains with (+) or without (-) the chrIV/chrXIV translocation analyzed on YPD and YPD with different concentrations of nocodazole (YPD+NDZ).

One strain, JDY528, which gained tolerance to nocodazole, was selected for further analysis. Whole-genome sequencing was conducted using the Oxford Nanopore MinION platform. Genome assembly using Canu and correction using Nanopolish enabled comparison with the reference genome and identification of two loxPsym-mediated genome arrangements in the JDY528 strain. A translocation was observed between chromosomes IV and XIV (mediated by IV-5 and XIV-1), and a 484 kb duplication was found on chromosome IV (mediated by IV-7 and IV-9) (Fig. 3c). The presence of these chromosomal rearrangements was confirmed by junction PCR (Extended Data Fig. 6b).

Major causal mutations in response to such selective forces are expected to change gene location/context and may involve the formation of new chromosomal junctions that affect higher-order genome regulation^38^. Therefore, the 3D genome structure of JDY528 was assessed by Hi-C analysis to determine the impact of rearrangements on spatial location. Genome-wide Hi-C contact maps^39^ were compared between JDY528 and SparLox83. Clustering of centromeres and telomeres (Extended Data Fig. 6c, red dots on the map) was seen in both strains, as previously reported^40, 41^. Although one of the rearranged loxPsym sequences in JDY528 was near the centromere of chrIV (Extended Data Fig. 6d), the contact maps showed that the translocation and duplication events in JDY528 did not disrupt localization and clustering of centromeres and telomeres. The log2-ratio heatmap between JDY528 and SparLox83 was plotted^42^ to quantitatively compare the differences between the two strains. This analysis showed that contacts between chromosomes without genomic rearrangement in JDY528 and SparLox83 were similar (Fig. 3d). However, the duplication on chromosome IV led to elevated contacts with the duplicated region. The translocation between chromosome XIV and chromosome IV altered the contacts between these two chromosomes (Fig. 3d, black circle). Moreover, the contacts between chromosome IX and other chromosomes were weaker in JDY528 than in SparLox83, indicating a potential change in the nuclear space occupied by chromosome IX in JDY528 (Fig. 3d, black arrow).

To test whether the variation in chromosomal structure was sufficient to cause the observed nocodazole tolerance, a split-*URA3* reporter strategy^28^ was used to reconstruct the chrIV/chrXIV translocation of JDY528 in JDY524 (the initial strain lacking loxPsym sites, Fig. 3e). The strain carrying only this translocation grew better than the split-*URA3* inserted into JDY524 on medium with nocodazole (Fig. 3f), providing direct evidence that the chrIV/chrXIV translocation contributed to the observed nocodazole tolerance. These data indicate that loxPsym-mediated large-scale genomic rearrangements changed the 3D genome structure, generating an advantageous phenotype.

### Chromosomes harboring sparsely arranged loxPsym sites combined with synthetic yeast chromosomes can generate complicated structural variations

SCRaMbLE is widely used in directed evolution to acquire strains with distinctive traits ^24–26, 28, 30, 34^. Here, to boost SCRaMbLE with genome-wide large-scale rearrangements, SparLox83 chromosomes were combined with a Sc2.0 synthetic chromosome in a single strain. As a proof of concept, SparLox83 was mated with JDY541, the strain carrying synIII^33^, to generate JDY544. After five rounds of SCRaMbLE using ReSCuES without selective pressures, nine SCRaMbLEant colonies were randomly isolated and sequenced. The presence of loxPsym sites and PCRTags in synIII and chrIII allowed reads with loxPsym sites to be extracted and differentiated in SparLox83 and synIII.

First, SCRaMbLEd synIII was resolved in the sequenced strains. The arrangement of synIII in each strain was deconvoluted and visualized as described previously^18^ (Fig. 4a). Rearrangement types in the nine strains included inversions, deletions, and duplications. As an example, the pattern of rearrangements detected in JDY533 is illustrated as a dot-plot (Extended Data Fig. 7a). By comparing sequence order and orientation in all strains except JDY550, several commonly conserved sequences (orange) and non-conserved sequences (gray) were identified (Extended Data Fig. 7b), most of which were consistent with previously reported regions^22^. The regions at the ends of synIII were often deleted, but to different degrees in different strains (Extended Data Fig. 7b, light gray rectangles). Seven of the nine strains harbored a circular synIII, which was also observed in SCRaMbLEant cells previously^43^ (Fig. 4a, see asterisks). This circularization was confirmed by reads spanning the left and right ends of synIII. In addition, whole-chromosome duplications were identified in three strains (JDY550, JDY596, and JDY599).

**Fig. 4.**
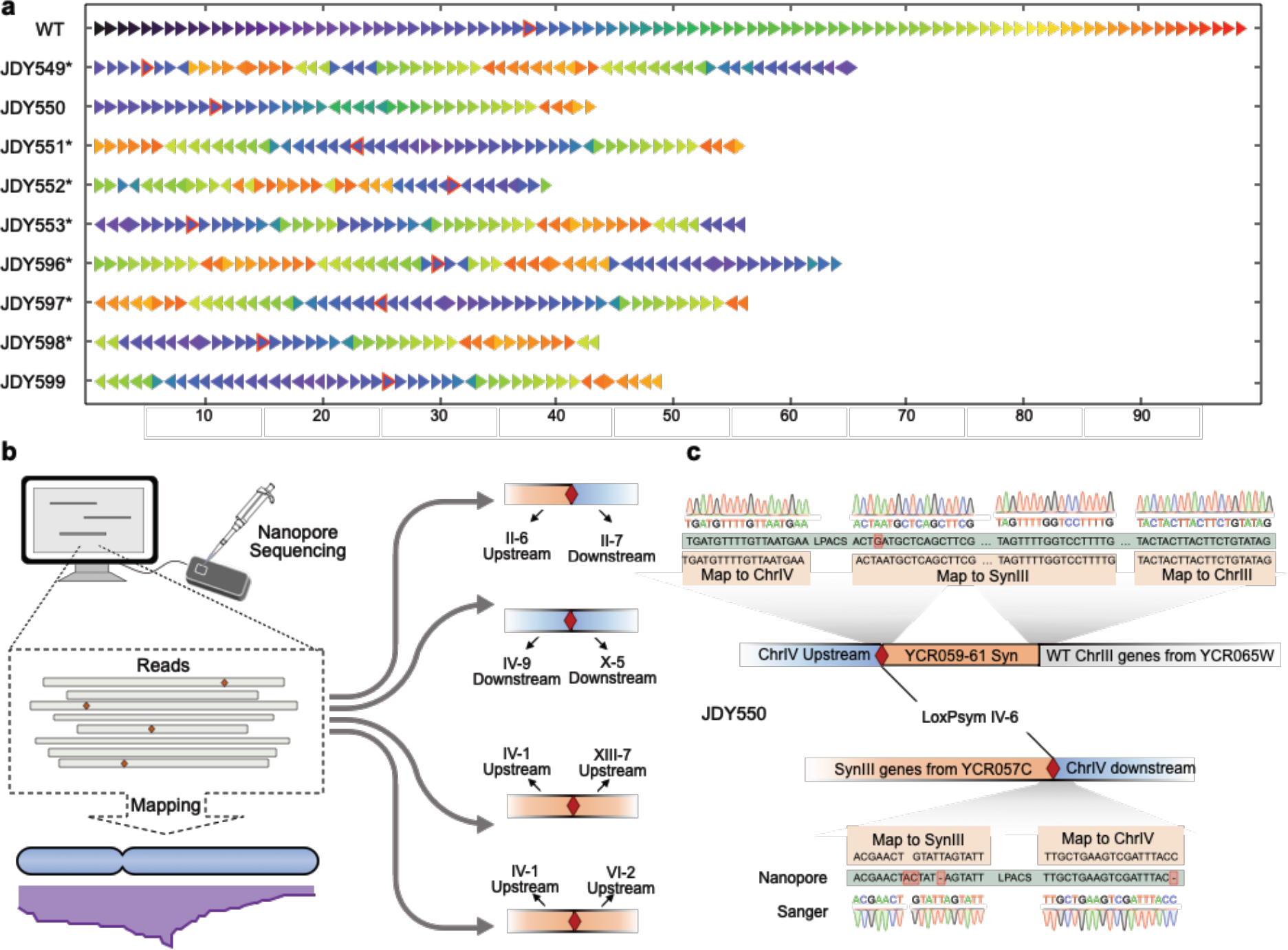
Structure variations caused by whole-genome-wide SCRaMbLE in diploid strains. **a**. SynIII rearrangements detected in diploids. Arrows indicate segments between two adjacent loxPsym sites. Arrow colors in the SCRaMbLEgram indicate the segment number in the parental chromosome, and arrow directions indicate orientation. Arrows with red borders denote segments containing centromeres. Strains with an asterisk (*) have a circular synIII as a result of SCRaMbLE. **b**. Schematic of Nanopore sequencing analysis of chromosome changes. A deletion between II-6 and II-7 was found in JDY550. A translocation between chrIV and chrX mediated by IV-9 and X-5 was found in JDY549, and the same loxPsym-mediated rearrangement also occurred in JDY598. A translocation between chrIV and chrXIII mediated by IV-1 and XIII-7 was found in JDY551, and the same loxPsym-mediated rearrangement also occurred in JDY597 and JDY599. A translocation between chrIV and chrVI mediated by IV-1 and VI-2 was found in JDY552. **c**. A complex rearrangement event among chromosomes III, IV, and synIII in JDY550. The translocation occurred at the IV-6 site and involved synIII and chrIII. This rearrangement was confirmed by both Nanopore and Sanger sequencing. Green bars show the sequence of one typical Nanopore read. LPACS, loxPsym and adjacent co-transformed sequence.

For chromosomes other than chromosome III and synIII, reads containing loxPsym sites were aligned to the reference sequences of SparLox83. Four types of structural variations were observed in seven of the nine strains, including one deletion event and six translocation events (Fig. 4b). In addition, a structural change was found in JDY550 mediated by IV-6 and loxPsym site 73, located between YCR057C and YCR059C on synIII. Read alignments revealed that IV-6-Up was combined with segment 74 on synIII, and IV-6-Down combined with segment 73 (Fig. 4c). A rearrangement between synIII and chrIII, in which the YCR061W gene was from synIII and its neighboring gene YCR065W was from chrIII (Fig. 4c), was also observed. However, no loxPsym sequence were detected at the breakpoint, which suggested that the rearrangement was generated by homologous recombination as synIII and chrIII shared significant sequence similarity. Three primer pairs were used to confirm these structure changes and the expected amplicons were directly sequenced to verify the specific translocation sites (Fig. 4c, Extended Data Figs. 7c&d). These results demonstrated that loxPsym sites in SparLox83 were able to combine with synthetic chromosomes to generate a range of structural variants genome wide, including rearrangements between SparLox83-derived loxPsym sites and synIII-derived loxPsym sites and one example of an homologous recombination-mediated rearragnment (Table S6).

### LOH and aneuploidy are frequently detected in SCRaMbLEd heterozygous diploids

As the rearrangement frequency was relatively lower with SparLox83-derived loxPsym sites than with synIII-based sites, it was doubtful whether LOH and aneuploidy, which were observed in previous SCRaMbLEd genomes^23^, were occurring in these strains. To examine this, the 2 kb upstream or downstream regions of each loxPsym site were designated as loxPsym site-Up and loxPsym site-Down and were analyzed as a unit. For example, X-1-Up and X-1-Down indicated the 2 kb upstream and downstream regions of X-1. Reads with or without loxPsym sites were mapped to the upstream/downstream regions and counted to deduce copy number variation of regions adjacent to loxPsym sites (Fig. 5a; where a blue square represents a copy of the upstream region or downstream region containing a loxPsym site and a white square represents a region lacking a loxPsym site). Using this strategy, copy numbers for all loxPsym upstream and downstream regions were assessed and some LOH events were observed in all nine strains (Fig. 5b). Several LOH events were likely caused by loxPsym site sequence deletion, such as the loxPsym sites on chromosome II (except for II-4 in JDY553 and JDY598), whereas other events were likely caused by loxPsym site sequence insertions into the counterpart of the other chromosome, such as IV-1 in all strains (except JDY552) (Fig. 5b). In addition, the structure of chrVIII in all nine strains differed from that of the parent strain JDY544. In JDY549, JDY550, and JDY 596, only one copy of wild-type chrVIII remained (Figs. 5b&c). In JDY551, JDY597, JDY598, and JDY 599, two copies of wild-type chrVIII were present instead of the heterozygous chrVIII in JDY544 (Figs. 5b&c). In JDY552, the loxPsym sequence of VIII-1 was inserted into the wild-type chrVIII and the remainder of SparLox83-based chrVIII was lost (Figs. 5b&c). JDY553 contained three copies of wild-type chrVIII, one of which contained an insertion of the VIII-1 loxPsym sequence (Figs. 5b&c). These variations were likely caused by structural differences in chrVIII between SparLox83 and the wild-type, in which a translocation of VIII-3 and IX-1 pre-existed prior to SCRaMbLEing (Extended Data Fig. 2).

**Fig. 5.**
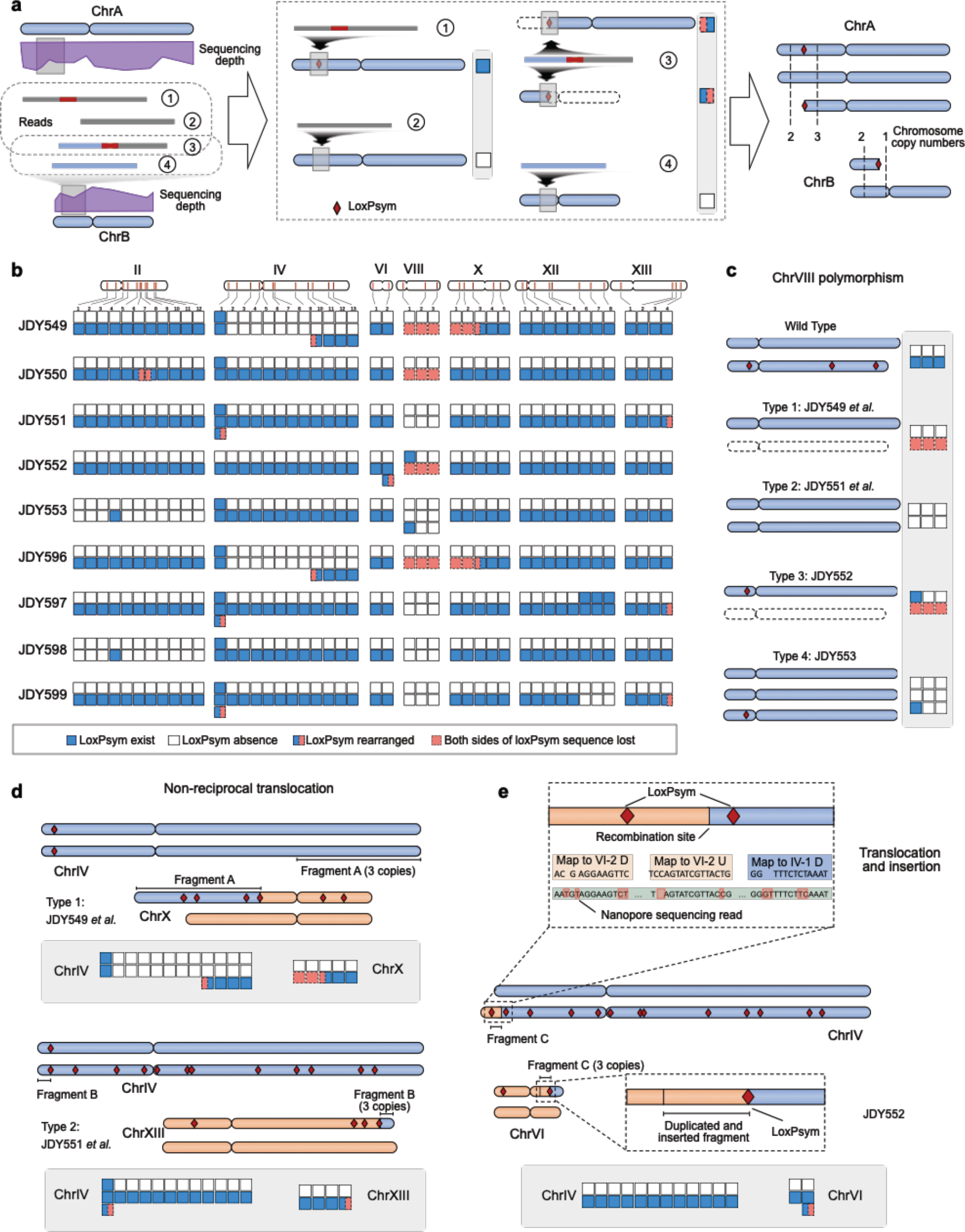
LOH and aneuploidy caused by genome-wide SCRaMbLE in diploids. **a**. Workflow of copy number variation analysis. Reads with or without loxPsym sites were mapped to the upstream/downstream regions of each loxPsym site. The copy number of the upstream/downstream regions for a certain loxPsym site were deduced based on read alignment. Different rectangle colors indicate four distinct situations deduced from read mapping: blue indicates loxPsym is present and not rearranged, white indicates absence of loxPsym, orange with dash lines indicates loss of loxPsym-containing sequencs and half bule with half orange squares indicate rearrangement. Structural variation was deduced using copy number calculation of loxPsym flanking sequences. **b**. Copy number variation of 2 kb upstream/downstream regions of the loxPsym sequence in the nine diploid strains. Chromosomes lacking rearrangements are not shown. **c**. Three different types of chrVIII polymorphism detected in the nine strains. **d**. Two types of non-reciprocal translocation observed between loxPsym sites IV-9 and X-5 or IV-1 and XIII-7. **e.** A translocation occurred between two loxPsym sites, IV-1 and VI-2, and a duplication of the region upstream of VI-2 was inserted at loxPsym site IV-1. Reads revealed that the upstream sequences (U) of VI-2 were connected to the downstream sequences (D) of IV-1. However, this was not mediated by loxPsym, indicating that a simultaneous duplication had occurred.

Using the copy number variation information, the chromosomal structural variations detected by WGS were reconstructed and several non-reciprocal translocations were identified. In JDY549 and JDY596, an extra copy of IV-9-Down combined with X-5-Down and formed a new chromosome, with deletion of X-5-Up (Fig. 4b and Fig. 5d). In JDY551, JDY597, and JDY599, an extra copy of IV-1-Up combined with XIII-7-Up, forming a new chromosome and deleting XIII-7-Down (Fig. 4b and Fig. 5d). In JDY552, however, a duplication of VI-2-Up was observed with no change to IV-1-Up (Fig. 5b). To clarify the events at these sites, reads were aligned to reference chromosomes. A translocation occurred between loxPsym sites IV-1 and VI-2, and a duplication of an approximately 45-kb region upstream of VI-2 was inserted upstream of IV-1. In this case, it is uncertain whether the rearrangement was loxPsym-mediated (Fig. 5e).

Overall, the diverse rearrangements and resulting genomes indicate that sparsely arranged loxPsym sites in SparLox83 can be used to generate combinatorial diversity with synthetic chromosomes, including structural variation, LOH, and aneuploidy.

### SparLox83 accelerates SCRaMbLE-mediated enhancement of acetic acid tolerance

The increased rearrangements observed in strains with synIII, as exemplified by the diversity of outcomes generated by SCRaMbLE of SparLox83 and synthetic chromosomes, suggested a potential strategy for strain improvement under stress conditions. To test this, the rate of variant evolution was compared between two heterozygous diploids, JDY546 (BY4741 × synIII) and JDY544 (SparLox83 × synIII), using tolerance to acetic acid (HAc) as a proof of principle. In total, 48 individual colonies from each strain were inoculated into a 96-well microplate and SCRaMbLE-Selection based evolution cycles were performed. Mixed strains after each SCRaMbLE round were selected on YPD plates with a HAc gradient (Fig. 6a).

**Fig. 6.**
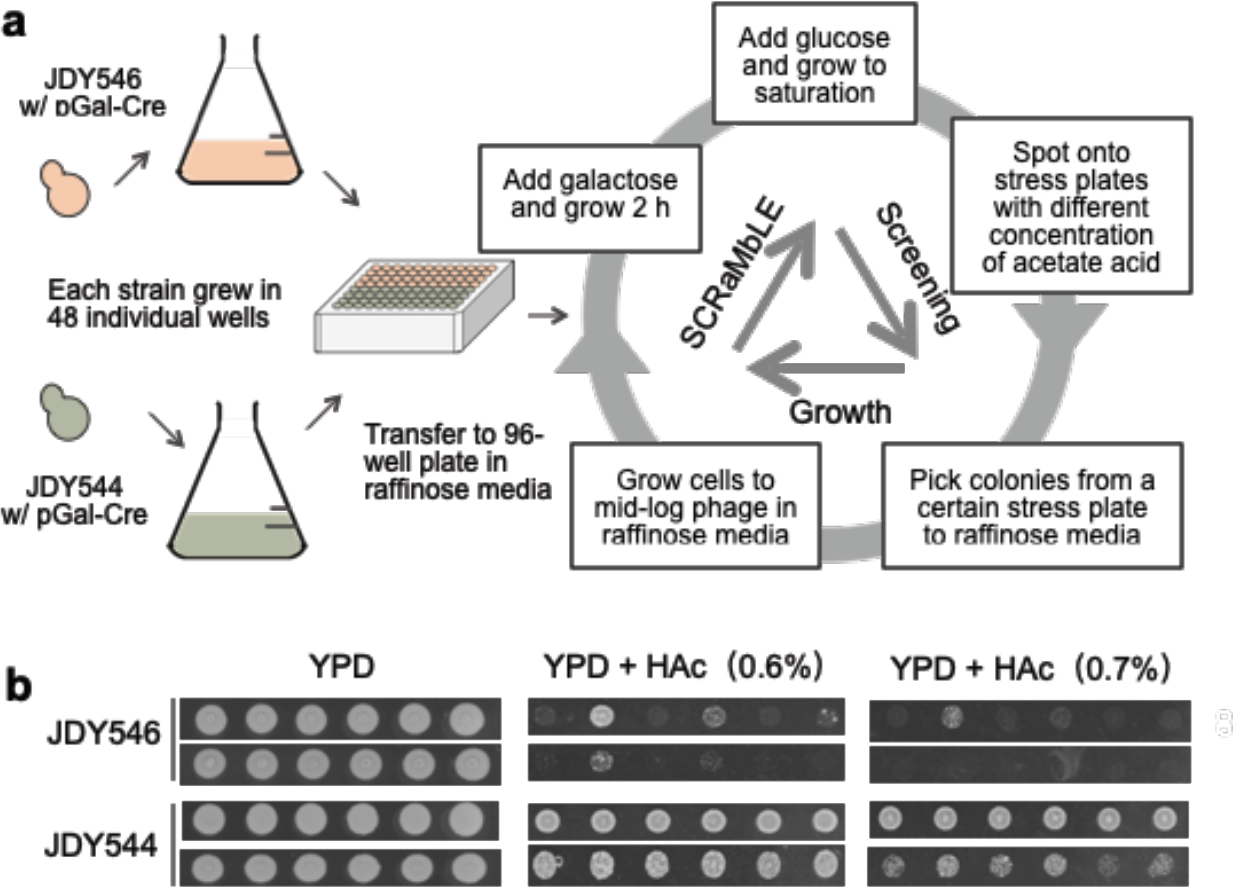
Rapid improvement of acetic acid tolerance using genome-wide SCRaMbLE. **a**. Workflow of SCRaMbLE-Selection based evolution cycles. Two heterozygous diploids, BY4741 × synIII (JDY546) and SparLox83 × synIII (JDY544), were inoculated into 48 individual wells and cycle growth-SCRaMbLE-screening was applied to SCRaMbLEant populations. Populations were screened after each round of SCRaMbLE by selection on YPD with an acetic acid (HAc) gradient. **b**. After three rounds of evolution, SCRaMbLEant populations from SparLox83 × synIII were more resistant to HAc than those from BY4741 × synIII.

The original strains (SparLox83, JDY544, and JDY546) were tested for HAc tolerance to identify an appropriate level of stress. Severe growth defects were seen in the presence of 0.4% HAc (Extended Data Fig. 8a), indicating that this concentration was appropriate for selection of tolerant SCRaMbLEants. After the first round of SCRaMbLE (R1), all populations were able to grow on media with a maximum of 0.4% HAc (Extended Data Fig. 8b). After the second (R2) and third (R3) rounds, the HAc-tolerance of all populations increased significantly (Extended Data Figs. 8b&c). After the second round, most SCRaMbLEant populations were able to grow on 0.5% HAc media, with populations from JDY544 exhibiting better growth than those from JDY546 (Extended Data Fig. 8b). After the third round (R3), two different concentrations of cells were tested on YPD plates with a gradient of HAc ranging from 0.5% to 0.8% (Fig. 6b and Extended Data Fig. 8c). The SCRaMbLEant populations from 48 individual JDY544 colonies were able to survive on 0.7% HAc, whereas only some of the 48 individual JDY546 colonies could grow under the same conditions. These results suggested that the development of increased HAc tolerance was greatly accelerated in SCRaMbLEant populations derived from JDY544 compared with those from JDY546, indicating that genomes with sparsely distributed loxPsym sites may be associated with rapid evolution of tolerance to HAc.

## Discussion

In this study, a strain containing 83 loxPsym sites sparsely arranged throughout all chromosome arms was constructed and used to study genomic rearrangement. Pooled sequencing of haploid and diploid SCRaMbLEants revealed that comparable inter- and intra-chromosomal rearrangement events were detected after inducing Cre recombinase expression. Our results indicated that genome ploidy and genomic location of loxPsym sites were the major factors that impacted rearrangement ability. Further examination revealed that one of the inter-chromosomal translocations altered the 3D genome structure and was associated with increased tolerance to environmental stress. When combined with SCRaMbLE in diploids made by crossing SparLox83 and a synIII-containing strain, LOH and aneuploidy, as well as structure variations, frequently occurred in the heterozygous diploid cells. The SparLox83-derived chromosomes in the diploids exhibited increased genome instability after Cre recombinase induction and generated diverse rearrangements to facilitate more rapid environmental adaptation than diploids only with wild-type chromosomes.

Genomic rearrangement is a major driver for rapid chromosomal evolution and the utility of the Cre-loxPsym system provides an artificial strategy to increase the opportunities for rearrangement^16^. Previous studies showed that several factors are associated with the recombination frequency mediated by the Cre-loxPsym system, including genome ploidy, chromosomal location of loxPsym sites, the spatial proximity of interactive loxPsym sites, and 3D chromosome conformation^18, 21, 31, 32^. Sequencing of pooled cells showed that loxPsym sites in SparLox83 exhibited uneven rearrangement event distribution and multiple rearrangement outliers were existed in both haploid and diploid strains. The intra-chromosomal outliers, mainly via the II-6 and II-7 loxPsym sites, were caused by higher rearrangement preference between these sites, which were only 7 kb apart and were therefore potentially situated in the same chromosomal interacting domain (CID)^44^. Supporting this notion, a previous study reported that loxPsym sites in the same CID actively rearrange with each other^21^. The mechanism of the Cre/loxP system relies on Cre recombinase functioning as a tetramer with two loxPsym sites, and our results are therefore consistent with the notion that the intra-chromosomal rearrangement frequency between two sites is correlated to their proximity^32^. However, no correlation was detected between proximity and inter-chromosomal rearrangement frequency^45^. Inter-chromosomal rearrangement may form dicentric chromosomes which can result in cell growth defect because dicentric chromosomes are instability at mitosis. Thus, the factors correlated to inter-chromosomal rearrangement are more complex than those of intra-chromosomal rearrangement.

Characterization of genetic rearrangements in SparLox83 and SparLox83 × BY4742 revealed that the proportion of rearrangement events was 10-fold higher in haploid than in diploid cells. There are two possible reasons behind this phenomenon. First, in haploid strains, loxPsym-mediated deletion of fragments might lead to lethality because of essential gene loss^46, 47^. However, deletions can occur in the diploid without lethal effects because of the additional presence of the wild-type genome^34^, thereby leading to a decreased total number of loxPsym sites. Second, the WT copy in diploid cells also led to LOH and aneuploidy during Cre expression, and this might decrease the total number of loxPsym sites (Fig. 4 and 5). In addition, five rounds of induction-selection were performed before sequencing, and this may have further lowered the total number of loxPsym sites.

Our introduction of SCRaMbLE in SparLox83 × synIII is the first trial to induce large-scale genomic rearrangement as well as SCRaMbLE in a diploid strain. Consistent with previous research^18^, the arrangements of synIII in nine diploid SCRaMbLEants obtained using ReSCuES showed unique structures that differed from one another (Fig. 4a), demonstrating the capacity of SCRaMbLE. Moreover, several synIII arrangement traits were found in these strains. First, the remaining segments showed similar biases in all strains (Extended Data Fig. 7b), partly overlapping earlier research^22^. Segments 65, 66, and 67 were deleted in our strains but were not retained in the haploid strain possessing an essential gene plasmid, indicating that synthetic lethal interactions can be bypassed when a copy of wild type chrIII is present. Second, all diploid SCRaMbLEants harbored a shorter synIII, whereas about one third of the haploid strains in previous studies harbored a rearranged synIII larger than the parental chromosome^18, 19^. These genotypic differences could be due to our use of linear chromosomes in a diploid strain, by contrast with previous studies that used ring chromosomes in haploid strains^18, 19^. These results indicate that, as previously noted^33^, deletion is more likely in diploid strains and large fragment deletions can cause severe growth defects in haploid strains. In addition, we found that SCRaMbLE generated a high frequency of chromosome circularization, with numerous segments near to the telomere region deleted but with gene arrangements near the centromere region retained.

Only one rearrangement between the SparLox83 and synIII chromosomes was observed in the nine diploid strains (Fig. 4c). However, a higher rate of intra-structural variations within synIII was identified (Fig. 4a). This could be caused by several potential mechanisms. First, synIII has a higher density of loxPsym sites and the sites are closer to one another than sites on the SparLox83 chromosomes, likely resulting in increased interactions between Cre recombinases binding at neighboring sites. Second, SCRaMbLEant sequencing analysis revealed that intra-chromosomal ARR is significantly higher than inter-chromosomal ARR (Fig. 2d), and similar outcomes were also previously observed in strains harboring more than one synthetic chromosome^19^. Third, high efficiency of recombination in yeast may inhibit the rearrangement efficiency mediated by loxPsym sites in diploid cells because donors with long homologous arms are provided by wild-type chromosomes and many LOH events are identified (Fig. 5).

In summary, we used a strategy of random incorporation of loxPsym sites across all yeast chromosomes to develop SparLox83, a strain with 83 loxPsym sites, and demonstrated the advantages of rapid environmental adaptation using the SCRaMbLE system. When construction of the final Sc2.0 yeast strains harboring synthetic chromosomes is complete and used with SparLox83, an increased frequency of genomic rearrangements between SparLox83-derived chromosomes and synthetic chromosomes can be expected in the hetero-diploid strain. In addition, more loxPsym sites can be incorporated into SparLox83, further increasing its rearrangement capacity and improving its utility in a broad range of applications.

## Methods

### Consecutive insertion of loxPsym sites

SparLox83 was generated using CRISPR/Cas9-mediated multiplex DNA integration as previously described^48^. Three donor-gRNA cassettes were amplified and co-transformed into yeast cells alongside linearized plasmid (200 ng each) using the lithium acetate transformation method. Two plasmids were used, containing either *LEU2* or *URA3*. Insertions were verified by PCR. Colonies containing repaired donor-gRNA-containing plasmids were eliminated by streaking onto YPGal plates (2% bacto-peptone, 1% yeast extract, 2% galactose and 2% agar). Colonies were subsequently streaked onto SC–Ura, SC–Leu and YPD plates to confirm loss of plasmid. Verified strains were used for the next round of loxPsym insertion.

### Yeast colony PCR

Cells were resuspended in 20 µL of 20 mM NaOH and lysed in a thermocycler (94 °C, 3 min; 4 °C, 2 min, 5 cycles). Cell lysate (1 µL) was used as template for PCR using a site-specific forward primer upstream of each insertion site and a common reverse primer, SHO582 (Table S4). PCR was performed using the following conditions: 94 °C for 5 min; 30 cycles of 94 °C, 30 s; 55 °C, 30 s; 72 °C, 1 min; and final extension at 72 °C for 7 min. Agarose gel analysis was used for visualization of PCR products.

### Preparation of Yeast Genomic DNA

Yeast genomic DNA was prepared as previously reported^49^ with minor modifications. Briefly, collected yeast cells were washed once in sterile water and resuspended in 100 µL breaking buffer (50 mM Tris pH8.0, 100 mM NaCl, 1% SDS, 2% Triton X-100, and 1 mM EDTA) before addition of an equal volume of glass beads (Sigma G8772) and 200 µL of Phenol-Chloroform-Isoamyl alcohol (PCI) (25:24:1). After vortexing for 10 min at room temperature, 100 µL sterile water was added and mixed by inverting the tube several times. After centrifugation at 12000 × g for 10 min, the upper layer (150 µL) was transferred to a new microfuge tube. For DNA precipitation, 500 µL 100% ethanol was added and the sample was incubated at -20 °C for 20 min. Genomic DNA was pelleted at 12000 × g for 5 min at 4 °C. The DNA pellet was dried at 45 °C for 15 min in vacuum centrifugal concentrator (Eppendorf Concentrator Plus) and then resuspended in 50 µL sterile water.

### Serial dilution assay

Yeast cells were cultured overnight in YPD medium with rotation at 30 °C. Cultures were adjusted to the same cell concentrations after measuring the optical density at 600 nm (OD_600_) and then serially diluted 10-fold with water. Diluted cells were spotted onto selective media plates, incubated at 30 °C for 48–72 hr, and then imaged for analysis.

### Assay for genomic stability

Yeast colonies were inoculated into 5 mL YPD medium and incubated at 30 °C for 24 hr with agitation. Cultures (5 μL) were transferred into new vessels containing 5 mL fresh YPD medium and cultured under the same conditions. These steps were repeated daily for ten days (approximately 120 generations), after which cells were streaked onto YPD plates and incubated at 30 °C for 1–2 days until colonies could be visualized. Single clones were isolated and subjected to PCR analysis to examine the presence of loxPsym sequences. In total, 35 independent clones were tested at 24 randomly chosen loxPsym sites.

### Pulsed-Field Gel Electrophoresis

Chromosomal DNA was prepared as previously described^50^. Samples were analyzed using a 0.9% agarose gel with 0.5× TBE (pH8.0) for 20 hrs at 14 °C on a BioRad CHEF Mapper apparatus. Voltage was 6 V/cm, at an angle of 120°, with 10–60 s switch time.

### SCRaMbLE of yeast cultures

SparL83 or the heterozygous diploids with ReSCuES sequence integrated were transformed with pGAL-Cre plasmid (clonNAT resistance). Single yeast colonies were inoculated into 5 mL YPD medium containing 0.1 mg/mL clonNAT and cultured overnight at 30 °C. Cultures were diluted to OD600 = 0.3 in 5 mL fresh YEP medium with 2% w/v raffinose and 0.1% w/v glucose and cultivated at 30 °C for 6 hr. Galactose was added to a final concentration of 2% w/v and cultures incubated at 30 °C for 2 h. Cultures were collected and plated onto solid YPD medium containing a selective agent.

For consecutive induction, five rounds of SCRaMbLE were induced as follows. SCRaMbLE was induced by the addition of galacotose as described above. Cells were collected and washed once with 1 mL sterile water, resuspended in 5 mL SC-Leu or SC-Ura medium in alternative rounds, and cultivated to the stationary phase.

### Split-*URA3* mediated rearrangment

The *URA3* gene was splitted in half by the intron of *ACT1*. A loxP sequence was inserted into the the intron of *ACT1*. To integrate the two halves of the URA3 gene into the genome, a His marker was fused to one half of the URA3 gene and a Leu marker was fused to another half by PCR. Using transformation-associated recombination, the two halves of the URA3 gene with different markers were inserted into the location of XIV-1 and IV-5 of JDY524, respectively. A colony growing on the SC-His-Leu palte were picked and transformed with pGAL-Cre plasmid. The Cre expression was inducted by the addition of galacotose as described above. Put the inducted culture onto the SC-Ura plate and incubate at 30 °C for 48 hrs. Colonies were verified by PCR.

### Screening of HAc tolerance strains by continuous evolution

JDY544 and JDY546 were transformed with pGAL-Cre plasmid (clonNAT resistance) and several rounds of SCRaMbLE-assisted evolution were performed as follows. Each strain was evolved in 48 individual populations. Single colonies were cultured overnight in YPD containing 0.1 mg/mL clonNAT at 30 °C and re-inoculated to obtain an OD600 = 0.3 in YEP medium with 2% w/v raffinose and 0.1% w/v glucose. After growing for 6 h at 30 °C, galactose (2% w/v final concentration) was added and cultures incubated at 30 °C for an additional 2 h to induce Cre expression. To inhibit Cre expression, glucose (2% w/v final concentration) was added and cultures incubated at 30 °C until saturated. The SCRaMbLEd populations were diluted 1:10 and 1:100 into 100 μL fresh YPD. Diluted cells were plated on YPD agar plates and stress plates with series concentration of HAc. The strains incubated at 30 °C for several days until visible colonies were formed for all the 96 populations on stress plates with low concentration of HAc. For the next round of evolution, the parent strains were picked from the stress plate with visible colonies for all the 96 populations. Repeat the incubation and induction processes.

### Whole-genome Nanopore sequencing

DNA extraction and sequencing were performed as described previously^29^. Briefly, genomic DNA was extracted according using a modified Qiagen Genomic-tip 100/g protocol with the Qiagen Genomic Buffer kit. DNA quality was assessed with agarose gel electrophoresis and with a NanoDrop™ 2000 Spectrophotometer and a Qubit 4 Fluorometer with dsDNA HS reagent. Oxford Nanopore Technology (ONT) sequencing library preparation was performed using SQK LSK108 or LSK109 kits with Native Barcoding kits EXP-NDB104 or EXP-NBD114, according to the manufacturer’s guidelines. Sequencing was performed on a MinION Mk1B device using FLO-MIN106D with R9.4.1 sequencing cells. Sequencing was performed for 48 hr using MinKNOW (v3.6.5) software.

### Junction detection of SCRaMbLEants

All possible combinations of the 83 loxPsym sites were assembled *in silico* using the yeast reference genome ((GCF_000146045.2). Only 1kb-long loxPsym neighboring sequences were retained. The assembled sequences were used as the reference for junction detection. LoxPsym-containing ONT sequence reads were identified using LAST software (v1250) and then aligned to the reference using Minimap2 (v2.20-r1061). Reads with MapQ > 20 that covered at least one reference sequence were retained for subsequent analysis. Reads containing more than one loxPsym site were analyzed separately. The Rearrangement Rate (RR) of a particular loxPsym site, i, was defined as Nrei/(2Nnormi+Nrei), where Nnormi and Nrei were the number of normal and rearranged reads for loxPsym site i. The Rearrangement Rate (RR) was subsequently normalized to the total number of rearranged loxPsym sites to calculate Average Rearrangement Rate (ARR). The Rearrangement Weight (RW) of two particular loxPsym sites, i and j, was calculated by equation 1, which also considered bias of sequencing depth.

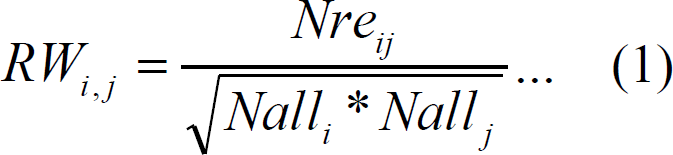

Nre_ij_ is the count of rearranged reads between loxPsym sites i and j, and Nalli = 2Nnormi+Nrei. Networks of inter-chromosomal and intra-chromosomal rearrangements were constructed using Cytoscape software (v3.7.1).

### Assembly-based structural variant calling

ONT reads (FASTQ format) were used with the Canu pipeling (v2.1.1) to assemble the SparLox83 and JDY528 genomes. Subsequently, Pilon software was employed to correct draft assemblies, utilizing the high-quality Illumina PE300 reads, for three consecutive rounds. The assembled genomes of JDY528 was aligned to SparLox83 using Minimap2 (v2.20-r1061) software with -x asm5 argument. The resultant PAF format files were used to display the structural variations.

### Structural and copy number variant and loss of heterozygosity detection in hetero-diploid strains

ONT reads (FASTQ format) from hetero-diploid strains were filtered and trimmed using NanoFilt software. LoxPsym-containing ONT reads were obtained by LAST software (v1250) for structural variant (SV) detection. To avoid alignment bias due to the similarities of ChrIII and SynIII, LAST was used to separate loxPsym-containing reads according to whether they contained SynIII PCRTags. Reads containing SynIII PCRTags were mapped to SynIII and were used to reconstruct SCRaMbLEd SynIII as previously described^18^. Reads without SynIII PCRTags were aligned to the SparLox83 genome for SV calling using the NGMLR-sniffles (v1.0.12 for sniffles and v0.2.7 for ngmlr) pipeline. In addition, SynIII PCRTag-containing reads were aligned to the SparLox83 genome to detect translocation between SynIII and SparLox83. LoxPsym sites that were not identified in any reads were considered to be the result of Loss of Heterozygosity (LOH). LoxPsym copy number variation (CNV) was identified using upstream and downstream 2000 bp flanking regions as described previously^20^.

### Hi-C library preparation and data processing

Hi-C library preparation was performed as previously described^51^ at Beijing Novogene Bioinformatics Technology Co. Ltd. Samples were flash-frozen and pulverized before formaldehyde crosslinking. Cell lysis and *Mbo*I digestion were performed after crosslinking, then DNA ends were labeled with biotin and joined. DNA was decrosslinked using Protease K and SDS. Free DNA was purified and extracted using Ampure XP beads. After quality control, ultrasonication was used to produce 200–500 bp DNA fragments. The biotin-labeled fragments were further enriched and sequenced by Illumina Novaseq. Hi-C data were mapped to the public S288C reference genome (GCF_000146045.2) using distiller-nf (v0.3.3). Read pairs that were not uniquely mapped (mapping score < 1) were discarded. Valid alignment files were transformed to .hic files for juicer using pre command. Hi-C map resolution was calculated as described previously^52^. Contact matrices used for heatmap visualization and further analysis were KR (Knight-Ruiz) normalized at 5 kb resolution. To compare the interaction frequencies between Spar83L and other strains, the log_2_ ratio of a 5 kb binned contact map was computed and normalized by subtracting the median of its values.

For the construction of a 3D chromosome model, Hi-C data were mapped to the SparLox83 genome and binned at 5 kb resolution. Chromosomal 3D structures were then inferred using the Pastis (v0.4) method with a Poisson model and visualized using PyMOL.

## Acknowledgements

This work was supported by the National Key Research and Development Program of China (2017YFA0205503), National Natural Science Foundation (21890743, 31725002, 32122050, 32101184), Youth Innovation Promotion Association CAS (2021359), Guangdong Natural Science Funds for Distinguished Young Scholar (2021B1515020060), Guangdong Provincial Key Laboratory of Synthetic Genomics (2019B030301006), Shenzhen Science and Technology Program (KQTD20180413181837372, 20200925153547003) and Shenzhen Outstanding Talents Training Fund. This work was also supported by UK Biotechnology and Biological Sciences Research Council (BBSRC) grants BB/M005690/1, BB/P02114X/1 and BB/W014483/1, Royal Society Newton Advanced Fellowship (NAF\R2\180590) and a Volkswagen Foundation “Life? Initiative” Grant (Ref. 94 771) to YC. WZ and JDB were supported by US NSF grants MCB-1026068, MCB-1443299, MCB-1616111 and MCB-1921641. We thank Gabrielle David for proof-reading this manuscript.

## Author contributions

J.D. conceived and designed the study. JD, CH, YM, YC supervised the experiments. TL, SZ, LC, SH, ZL, WY, SJ, MM, DS, WZ performed the experiments. JX, SZ and CH analyzed HiC data. JD, TL, SJ, LC, ZL, YM, YC, JDB analyzed the data and wrote the manuscript. All authors have read and approved the final manuscript.

## References

1. Perry, G. H. et al. Copy number variation and evolution in humans and chimpanzees. Genome Res. 18, 1698–1710 (2008).

2. Rieseberg, L. H. Chromosomal rearrangements and speciation. Trends Ecol. Evol. 16, 351–358 (2001).

3. Lee, J. A. & Lupski, J. R. Genomic Rearrangements and Gene Copy-Number Alterations as a Cause of Nervous System Disorders. Neuron 52, 103–121 (2006).

4. Lalani, S. R. Other genomic disorders and congenital heart disease. Am. J. Med. Genet. Part C Semin. Med. Genet. 184, 107–115 (2020).

5. Stephens, P. J. et al. Massive genomic rearrangement acquired in a single catastrophic event during cancer development. Cell 144, 27–40 (2011).

6. Dixon, J. R. et al. Integrative detection and analysis of structural variation in cancer genomes. Nat. Genet. 50, 1388–1398 (2018).

7. Ehman, E. C. et al. Identification of focally amplified lineage-specific super-enhancers in human epithelial cancers. Nat. Genet. 48, 176–182 (2016).

8. Hnisz, D. et al. Activation of proto-oncogenes by disruption of chromosome neighborhoods. Science (80-.). 351, 1454–1458 (2016).

9. Hurles, M. E., Dermitzakis, E. T. & Tyler-Smith, C. The functional impact of structural variation in humans Introduction to genomic structural variation. Trends Genet. 24, 238–245 (2010).

10. Weischenfeldt, J. et al. Pan-cancer analysis of somatic copy-number alterations implicates IRS4 and IGF2 in enhancer hijacking. Nat. Genet. 49, 65–74 (2017).

11. Lupski, J. R. Genomic rearrangements and sporadic disease. Nat. Genet. 39, S43–S46 (2007).

12. Periwal, V. & Scaria, V. Insights into structural variations and genome rearrangements in prokaryotic genomes. Bioinformatics 31, 1–9 (2015).

13. Deineri, D. et al. Engineering evolution to study speciation in yeasts. Nature 422, 68–72 (2003).

14. Fleiss, A. et al. Reshuffling yeast chromosomes with CRISPR Cas9. PLoS Genet. 15, 1–26 (2019).

15. Richardson, C. & Jasin, M. Frequent chromosomal translocations induced by DNA double-strand breaks. Nature 405, 697–700 (2000).

16. Dymond, J. S. et al. Synthetic chromosome arms function in yeast and generate phenotypic diversity by design. Nature 477, 471–476 (2011).

17. Mercy, G. et al. 3D organization of synthetic and scrambled chromosomes. Science (80-.). 355, eaaf4597 (2017).

18. Shen, Y. et al. SCRaMbLE generates designed combinatorial stochastic diversity in synthetic chromosomes. Genome Res. 26, 36–49 (2016).

19. Wang, J. et al. Ring synthetic chromosome V SCRaMbLE. Nat. Commun. 9, 1–9 (2018).

20. Shen, Y. et al. Deep functional analysis of synII, a 770-kilobase synthetic yeast chromosome. Science (80-.). 355, (2017).

21. Luo, Z. et al. Compacting a synthetic yeast chromosome arm. Genome Biol. 22, 1–18 (2021).

22. Wang, P. et al. Scrambleing of a synthetic yeast chromosome with clustered essential genes reveals synthetic lethal interactions. ACS Synth. Biol. 9, 1181– 1189 (2020).

23. Li, Y. et al. Loss of heterozygosity by SCRaMbLEing. Sci. China Life Sci. 62, 381–393 (2019).

24. Blount, B. A. et al. Rapid host strain improvement by in vivo rearrangement of a synthetic yeast chromosome. Nat. Commun. 9, (2018).

25. Jia, B. et al. Precise control of SCRaMbLE in synthetic haploid and diploid yeast. Nat. Commun. 9, (2018).

26. Luo, Z. et al. Identifying and characterizing SCRaMbLEd synthetic yeast using ReSCuES. Nat. Commun. 9, 1–10 (2018).

27. Liu, W. et al. Rapid pathway prototyping and engineering using in vitro and in vivo synthetic genome SCRaMbLE-in methods. Nat. Commun. 9, 1–12 (2018).

28. Wu, Y. et al. In vitro DNA SCRaMbLE. Nat. Commun. 9, (2018).

29. Ong, J. Y. et al. Scramble: A study of its robustness and challenges through enhancement of hygromycin b resistance in a semi-synthetic yeast. Bioengineering 8, 1–15 (2021).

30. Gowers, G. O. F. et al. Improved betulinic acid biosynthesis using synthetic yeast chromosome recombination and semi-automated rapid LC-MS screening. Nat. Commun. 11, (2020).

31. Zhang, H. et al. Systematic dissection of key factors governing recombination outcomes by GCE-SCRaMbLE. Nat. Commun. 13, 5836 (2022).

32. Zhou, S. et al. Dynamics of synthetic yeast chromosome evolution shaped by hierarchical chromatin organization. bioRxiv 2021.07.19.453002 (2021).

33. Annaluru, N. et al. Designer Eukaryotic Chromosome. Science 344, 55–59 (2014).

34. Shen, M. J. et al. Heterozygous diploid and interspecies SCRaMbLEing. Nat. Commun. 9, (2018).

35. Schep, A. N. et al. Structured nucleosome fingerprints enable high-resolution mapping of chromatin architecture within regulatory regions. Genome Res. 25, 1757–1770 (2015).

36. Chang, S. L., Lai, H. Y., Tung, S. Y. & Leu, J. Y. Dynamic Large-Scale Chromosomal Rearrangements Fuel Rapid Adaptation in Yeast Populations. PLoS Genet. 9, (2013).

37. Tang, X. X. et al. Origin, regulation, and fitness effect of chromosomal rearrangements in the yeast saccharomyces cerevisiae. Int. J. Mol. Sci. 22, 1–17 (2021).

38. Maass, P. G., Barutcu, A. R., Weiner, C. L. & Rinn, J. L. Inter-chromosomal Contact Properties in Live-Cell Imaging and in Hi-C. Mol. Cell 69, 1039–1045.e3 (2018).

39. Lieberman-Aiden, E. et al. Comprehensive Mapping of Long-Range Interactions Reveals Folding Principles of the Human Genome. Science (80-.). 326, 289–293 (2009).

40. Cournac, A., Marie-Nelly, H., Marbouty, M., Koszul, R. & Mozziconacci, J. Normalization of a chromosomal contact map. BMC Genomics 13, (2012).

41. Noble, W. et al. A Three-Dimensional Model of the Yeast Genome. in Lecture Notes in Computer Science (including subseries Lecture Notes in Artificial Intelligence and Lecture Notes in Bioinformatics) vol. 6577 LNBI 320–320 (2011).

42. Ruault, M. et al. Sir3 mediates long-range chromosome interactions in budding yeast. Genome Res. 31, 411–425 (2021).

43. Xiong, Y. et al. Structural Variations and Adaptations of Synthetic Chromosome Ends Driven by SCRaMbLE in Haploid and Diploid Yeasts. (2022) doi:10.1021/acssynbio.2c00424.

44. Hsieh, T. H. S. et al. Mapping Nucleosome Resolution Chromosome Folding in Yeast by Micro-C. Cell 162, 108–119 (2015).

45. Jager, D. & Philippsen, P. Stabilization of dicentric chromosomes in Saccharomyces cerevisiae by telomere addition to broken ends or by centromere deletion. EMBO Journal vol. 8 247–254 (1989).

46. Giaever, G. et al. Functional profiling of the Saccharomyces cerevisiae genome. Nature 418, 387–391 (2002).

47. Winzeler, E. A. et al. Functional characterization of the S. cerevisiae genome by gene deletion and parallel analysis. Science (80-.). 285, 901–906 (1999).

48. Hou, S., Qin, Q. & Dai, J. Wicket: A Versatile Tool for the Integration and Optimization of Exogenous Pathways in Saccharomyces cerevisiae. ACS Synth. Biol. 7, 782–788 (2018).

49. Zhang, W. et al. Engineering the ribosomal DNA in a megabase synthetic chromosome. Science 355, (2017).

50. Schwartz, D. C. & Cantor, C. R. Separation of yeast chromosome-sized DNAs by pulsed field gradient gel electrophoresis. Cell 37, 67–75 (1984).

51. Belton, J. M. et al. Hi-C: A comprehensive technique to capture the conformation of genomes. Methods 58, 268–276 (2012).

52. Rao, S. S. P. et al. A 3D map of the human genome at kilobase resolution reveals principles of chromatin looping. Cell 159, 1665–1680 (2014).

