## Supplemental figures and tables for "Large-scale genomic rearrangements boost SCRaMbLE in *Saccharomyces cerevisiae*"

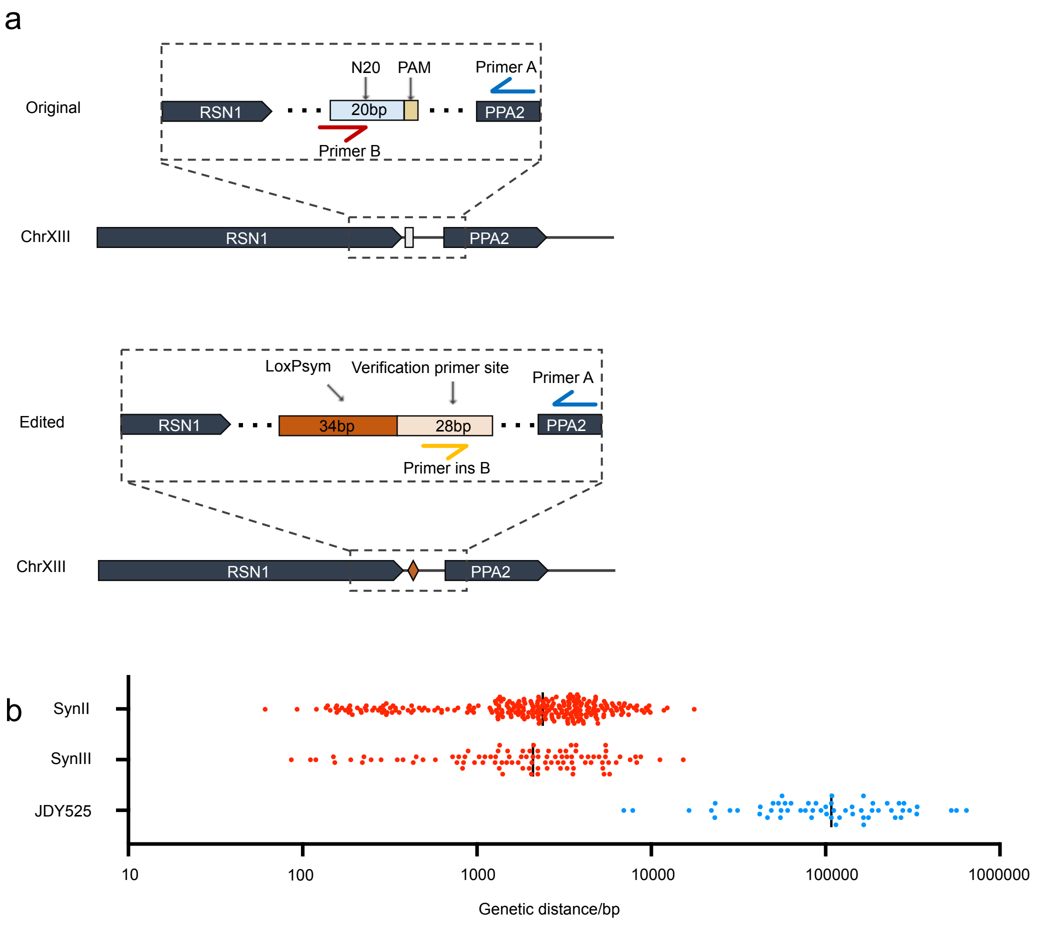


**Extended Data Fig. 1 | Selection of loxPsym insertion sites. a.** Example of loxPsym insertion site selection. Intergenic regions of the yeast genome were used as inputs to search for possible crRNA (protospacer) sequences for Cas9 nuclease (denoted N20) at http://crispr-era.stanford.edu/. Next, the secondary structure of gRNAs with the highest scores according to http://rna.tbi.univie.ac.at/cgi-bin/RNAfold.cgi were determined, and sequences lacking a hairpin and with lower minimum free energy structures were selected. For each loxPsym insertion event, primer A was designed within the target locus, primer B was designed adjacent to the sequence being targeted by the insertion event, and primer insB was designed specific to the unique sequence inserted with the loxPsym site. When loxPsym sites were inserted correctly (e.g., JDY525), primer pair A + B would not produce an ampliconfrom genomic DNA, but primer pair A + insB would produce amplicons of the expected sizes. **b.** Genetic distance between two adjacent loxPsym is different between synthetic chromosomes and JDY525. Each dot represents a pair of adjacent loxPsym sites.

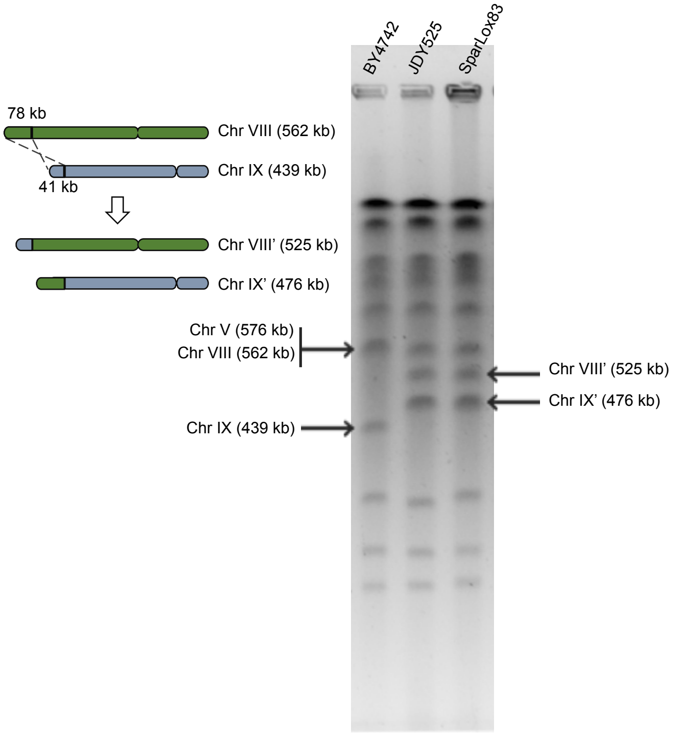


**Extended Data Fig. 2 | Translocation occurring during strain construction.** Pulsed-field gel electrophoresis analysis of a translocation between chromosomes VIII and IX in JDY525 and SparLox83. This translocation occurred between the inserted loxPsym sites VIII-3 and IX-1 and resulted in size alterations to both chromosomes (arrows).

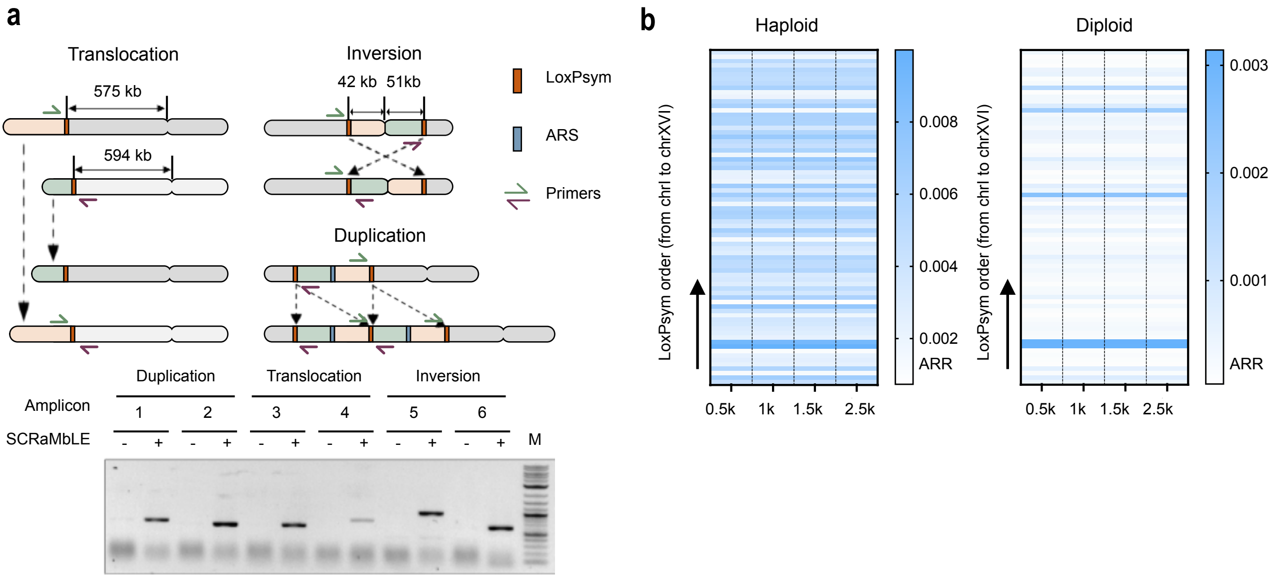


**Extended Data Fig. 3 | Junction PCR and Nanopore sequencing revealed diverse rearrangements in SparLox83 after SCRaMbLE. a**. Junction PCR to diagnose junctions formed from different types of rearrangement. Examples of the following are shown: translocation, with two loxPsym sites on different chromosomes at similar distances from their original centromeres; inversion, with two loxPsym sites on different arms of the same chromosome at similar distances from the centromere; and duplication, with the fragment between two loxPsym sites containing at least one autonomously replicating sequence (ARS). Primer pairs were designed to generate 500–1000 bp amplicons only upon rearrangement. Amplicon 1, duplication mediated by XIII-5 and XIII-6; amplicon 2, duplication mediated by IV-1 and IV-2; amplicon 3, translocation mediated by IV-9 and XII-6; amplicon 4, translocation mediated by VII-2 and X-4; amplicon 5, inversion mediated by X-5 and X-6; and amplicon 6, inversion mediated by I-2 and I-3. Genomic template DNA was from SparLox83 (-) or the rearrangement induction population (+). **b**. ARR calculated using different flank region lengths in haploid and diploid cells. X-axis, length of flank regions; Y-axis, loxPsym sites arranged in order.

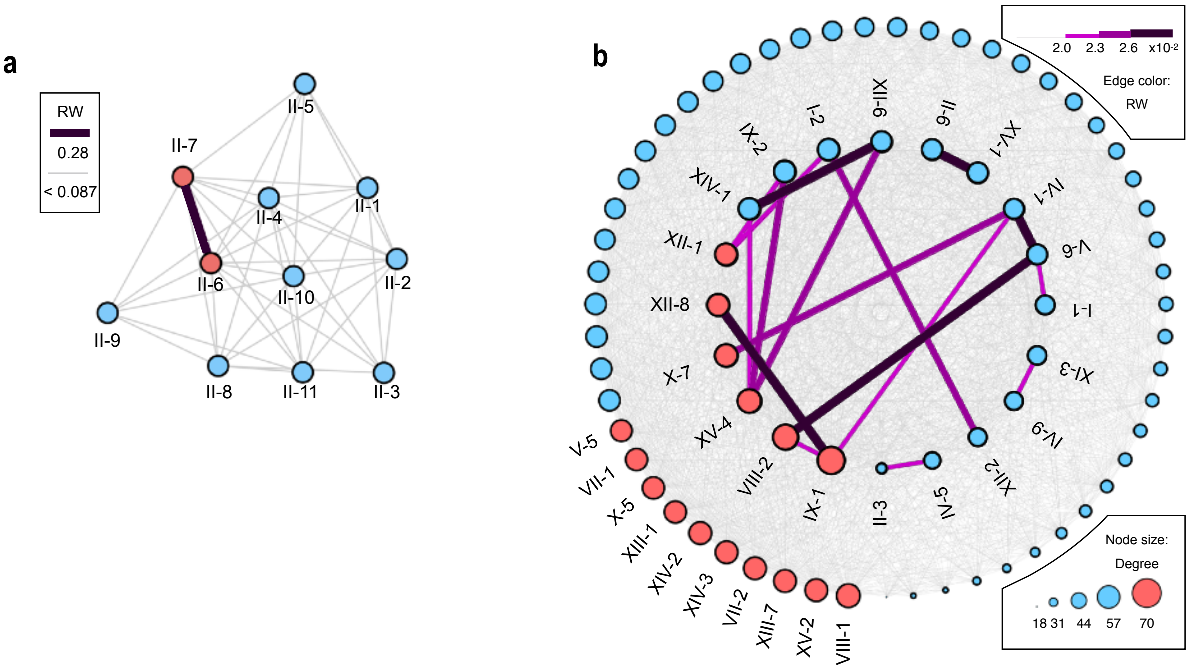


**Extended Data Fig. 4 | Uneven distribution of rearrangement frequencies at different loxP sites.** **a**. Intrachromosomal rearrangement network for chrII, representing possible rearrangements among the eleven loxPsym sites (nodes). Pairs of loxPsym sites are shown connected if a rearrangement was detected. Line thickness is proportional to the number of rearrangements detected. The network was generated using Cytoscape (v3.7.1). **b**. Interchromosomal rearrangement network in haploid cells. Node size indicates the number of interconnections with other sites (degree). Red dots indicate degree >= 60. The internal circle consists of dots with rearrangement weights (RW) > 2.0 × $10^{-2}$.

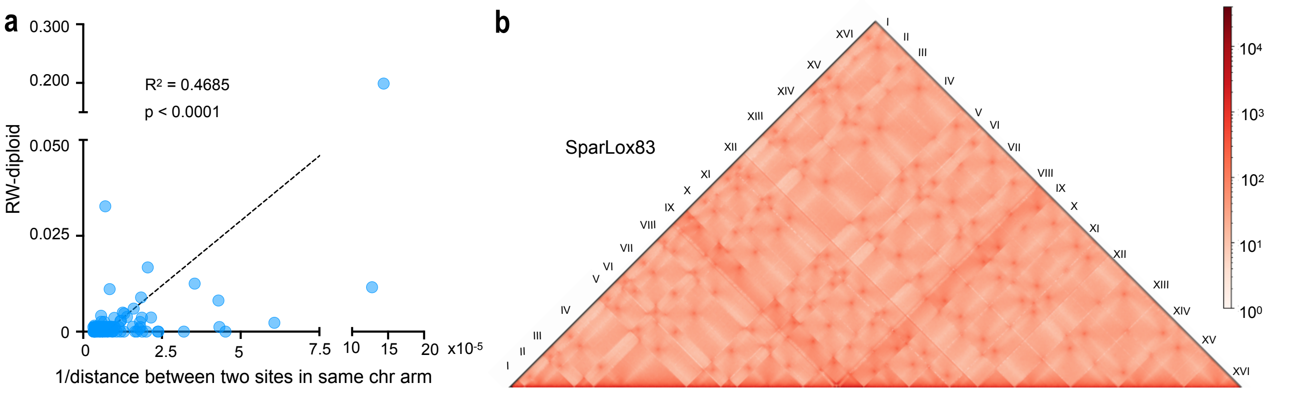


**Extended Data Fig. 5 | Correlation between genomic location or 3D spatial distancing of loxPsym sites and their rearrangement event frequencies. a.** Correlation between RW in diploid cells and 1/genomic distance between loxPsym sites in the same chromosome arm. $R^2$=0.4685 and p<0.0001. **b.** KR-normalized Hi-C contact map of SparLox83.

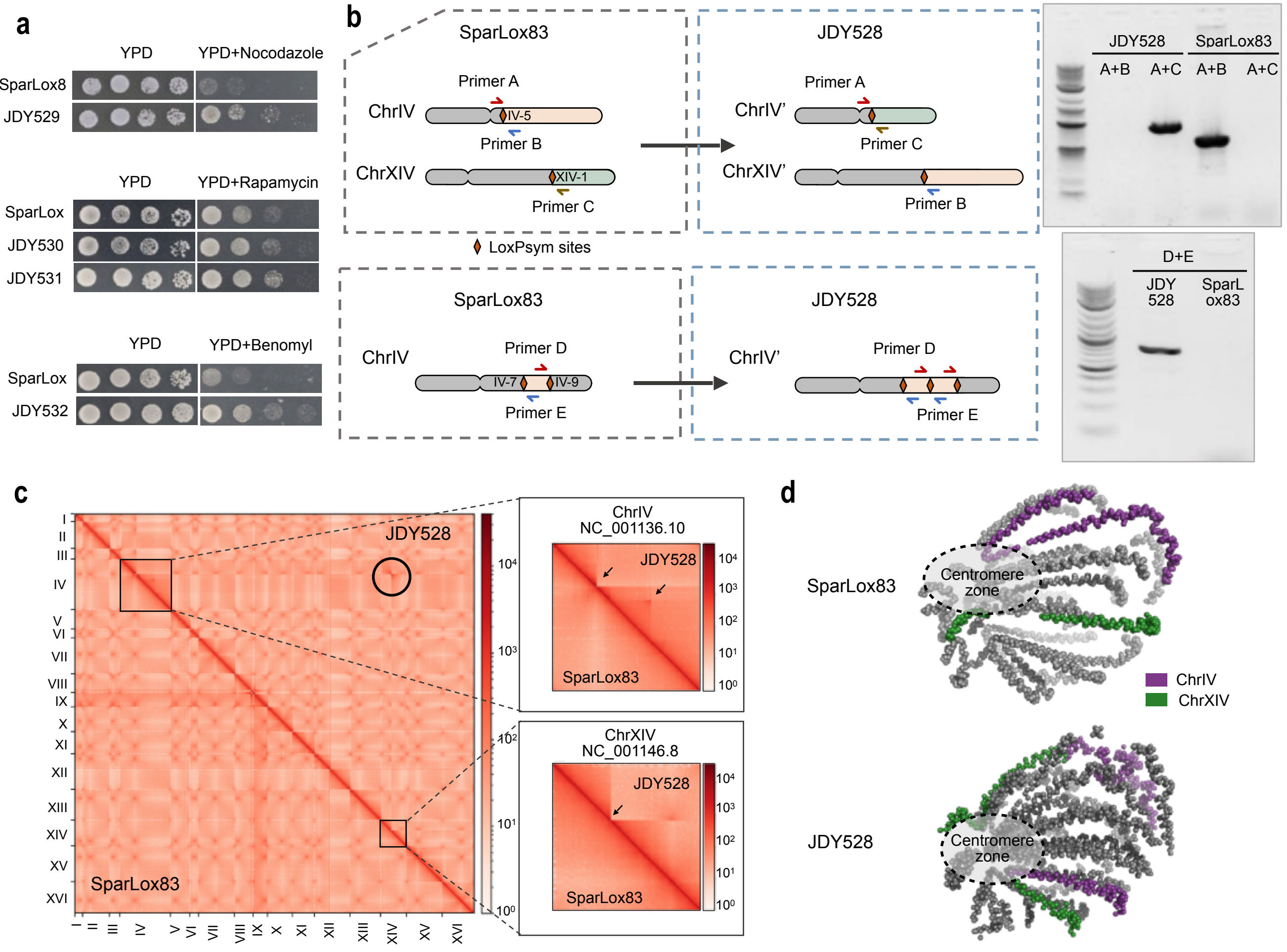


**Extended Data Fig. 6 | Exploration of the impact of large-scale rearrangement on phenotype, transcription, and genome 3D structure upon exposure to stress. a**. Stress tolerance of selected strains after SCRaMbLE. Strains at log phase were 10-fold serially diluted onto YPD agar plates containing 10 μg/mL nocodazole, 40 μg/mL benomyl, or 10 ng/mL rapamycin and incubated for 2 days at 30°C. **b**. PCR verification of rearrangements in JDY528. Primers are in Table S4. **c**. Contact maps of the SparLox83 and JDY528 genomes. Translocations and duplications are indicated with black arrows. Red dots in the map indicate centromeres. **d**. Predicted average 3D chromosome localization in SparLox83 and JDY528 based on contact mapping. Rearranged chromosomes are highlighted in different colors.

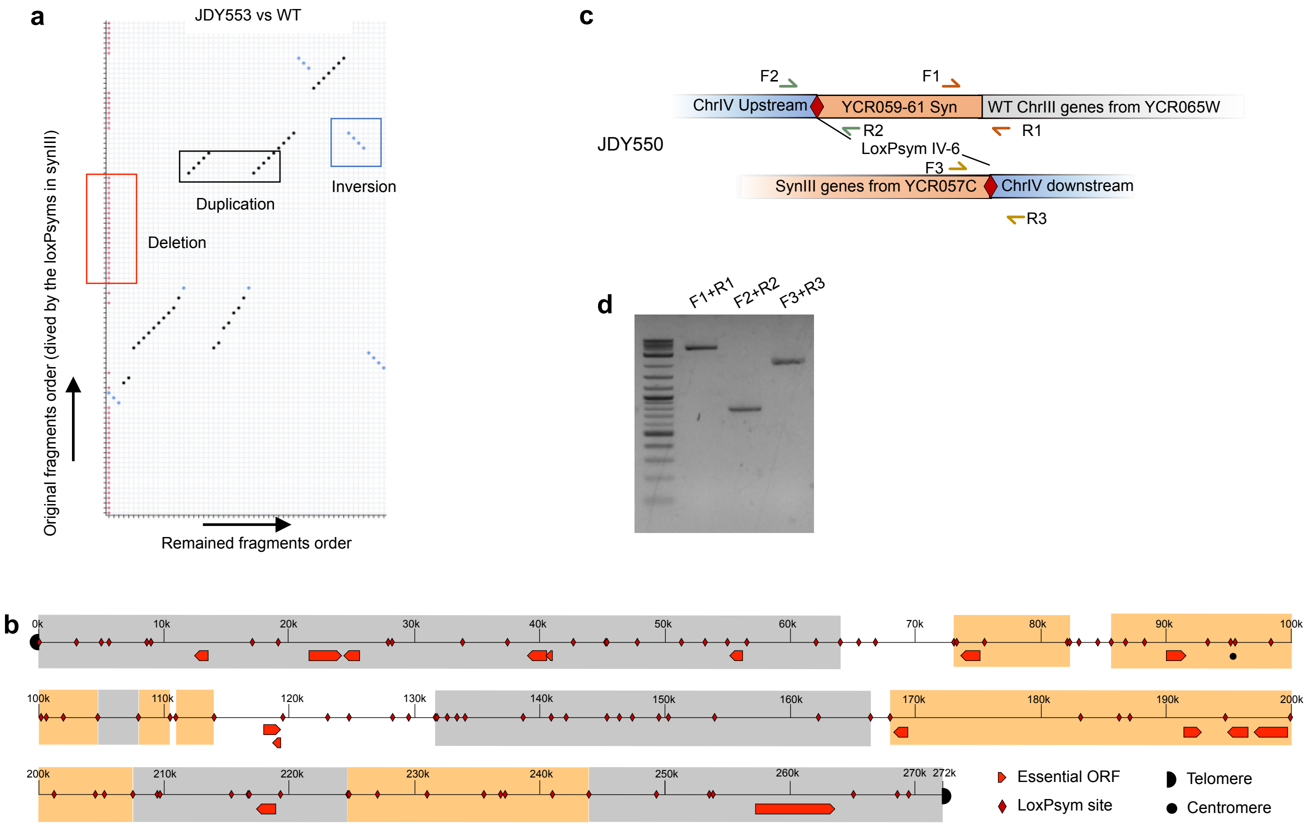


**Extended Data Fig. 7 | Consecutive whole-genome-wide SCRaMbLE in other diploid strains. a**. Dot-plots illustrating synIII rearrangements in JDY553. Each dot represents a fragment between two loxPsym sites. Black dots represent fragments arranged in synIII sequence order and blue dots represent fragments inverted with respect to the synIII sequence. **b**. Schematic representation of synIII illustrating common rearrangements among SCRaMbLEants. Regions highlighted in orange represent sequences retained after SCRaMbLE in all SCRaMbLEants. Regions highlighted in gray were lost from synIII in all SCRaMbLEants. Red arrows indicate open reading frames (ORFs) of essential genes. **c**. PCR primers designed to verify translocation between SparLox83 and synIII chromosomes in strain JDY550. **D**. PCR analysis of JDY550. Amplicons were sequenced and the data are shown in Fig 4c.

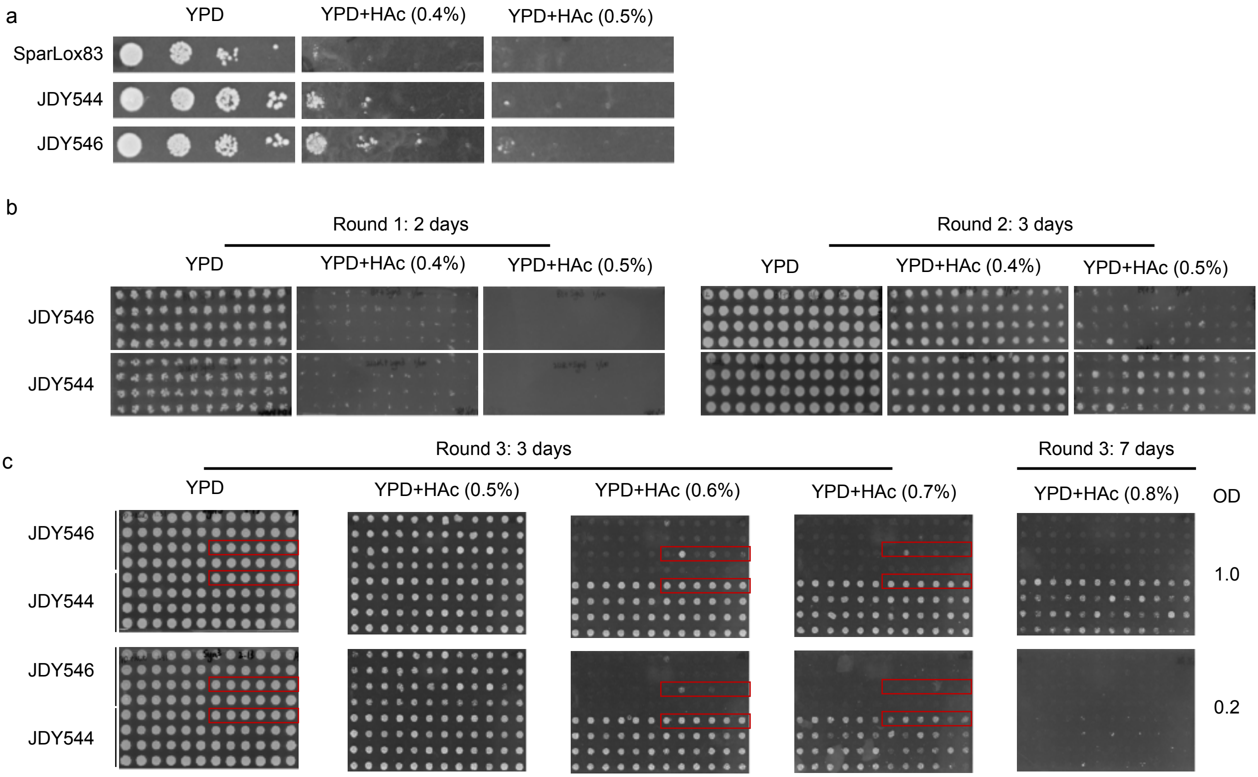


**Extended Data Fig. 8 | Rapid development of HAc tolerance using whole-genome-wide SCRaMbLE in the presence of synIII or native chrIII. a**. Serial dilution assay (10-fold) on YPD medium with/without HAc. Three strains, SparLox83, BY4741 × synIII, and SparLox83 × synIII, all showed the expected growth defect on YPD with 0.4% and 0.5% HAc. **b**. Growth of 96 SCRaMbLEant populations after first and second round SCRaMbLE. After SCRaMbLE, the saturated culture was diluted 10- or 100-fold and spotted onto YPD with/without HAc. **c**. Growth of 96 populations after third round SCRaMbLE. After SCRaMbLE, a 96-well plate reader was used to measure the density of all 96 populations. $A_{600}$ values were adjusted to 1.0 and 0.2 and cells were spotted onto plates. Red boxes indicate the colonies shown in Fig. 6b.

### Supplementary Tables

#### Table S1. Locations of inserted loxPsym sites in SparLox83

| Name | Insert site[a] | Notes | Name | Insert site[a] | Notes |
|---|---|---|---|---|---|
| I-1 | 51708 | | VIII-1 | 74959 | |
| I-2 | 115050 | | VIII-2 | 345978 | |
| I-3 | 209083 | | VIII-3 | 489110 | |
| II-1 | 56346 | | IX-1 | 77997 | |
| II-2 | 176659 | | IX-2 | 411714 | Self-duplication[b] |
| II-3 | 253209 | Off-target | IX-3 | 413059 | |
| II-4 | 325134 | | X-1 | 130456 | Self-duplication |
| II-5 | 433039 | | X-2* | 131410 | |
| II-6 | 479449 | | X-3 | 188979 | Self-duplication |
| II-7 | 486400 | | X-4 | 190016 | |
| II-8 | 541862 | Off-target | X-5 | 303740 | |
| II-9 | 563996 | | X-6 | 554414 | |
| II-10 | 648545 | | X-7 | 642488 | |
| II-11 | 671827 | | XI-1 | 74611 | |
| III-1 | 24306 | | XI-2 | 356299 | |
| III-2 | 200486 | | XI-3 | 582098 | |
| III-3 | 309197 | | XI-4 | 635781 | |
| IV-1 | 43937 | | XII-1 | 83403 | |
| IV-2 | 144091 | | XII-2 | 133362 | |
| IV-3 | 309055 | | XII-3 | 234581 | |
| IV-4 | 417044 | | XII-4 | 419248 | |
| IV-5 | 458920 | | XII-5 | 447506 | |
| IV-6 | 589165 | | XII-6 | 782897 | |
| IV-7 | 605638 | | XII-7 | 924842 | |
| IV-8 | 869600 | Off-target | XII-8 | 1090211 | |
| IV-9 | 1029771 | | XIII-1 | 119886 | Self-duplication |
| IV-10 | 1086043 | | XIII-2 | 120927 | |
| IV-11 | 1286916 | | XIII-3 | 121977 | |
| IV-12 | 1342001 | Self-duplication | XIII-4 | 123019 | |
| IV-13 | 1342956 | | XIII-5 | 762757 | |
| V-1 | 70795 | | XIII-6 | 804923 | |
| V-2 | 78619 | Off-target | XIII-7 | 865447 | |
| V-3 | 190095 | | XIV-1 | 295627 | |
| V-4 | 213214 | | XIV-2 | 572843 | |
| V-5 | 244493 | | XIV-3 | 656031 | |
| V-6 | 547928 | | XV-1 | 216750 | |
| VI-1 | 31506 | | XV-2 | 743244 | Self-duplication |

| | | | |
|---|---|---|---|
| VI-2 | 218228 | XV-3 | 743745 |
| VII-1 | 196662 | XV-4 | 908241 |
| VII-2 | 245412 | XVI-1 | 451392 |
| VII-3 | 325056 | XVI-2 | 689950 |
| VII-4 | 888555 | | |

[a]Sequence coordinates are from the Saccharomyces Genome Database.

[b]Self-duplication: more than one loxPsym sequence separated by vector backbone.

*The X-2 loxPsym sequence has one mismatch.

**Table S2. List of strains used in this study**

| Systematic Name | Mating type | Genotype | Description |
|---|---|---|---|
| JDY406 | a | *leu2Δ0 met15Δ0 ura3Δ0 his3Δ1* | BY4741 |
| JDY407 | alpha | *leu2Δ0 lys2Δ0 ura3Δ0 his3Δ1* | BY4742 |
| JDY408 | a/alpha | *his3Δ1/his3Δ1 leu2Δ0/leu2Δ0 LYS2/lys2Δ0 met15Δ0/MET15 ura3Δ0/ura3Δ0* | BY4743 |
| JDY524 | a | *his3Δ200 leu2Δ0 lys2Δ0 trp1Δ63 ura3Δ0 met15Δ0 ho:: pTDH3-Cas9-tCYC1-TRP1 can1::PED* | Initial strain, with Cas9 gene inserted into *HO* locus. |
| JDY525 | a | *his3Δ200 leu2Δ0 lys2Δ0 trp1Δ63 ura3Δ0 met15Δ0 ho:: pTDH3-Cas9-tCYC1-TRP1 can1::PED* | Final strain, with 83 loxPsym sites inserted across all 16 chromosomes. |
| JDY526 | a | *his3Δ200 leu2Δ0 lys2Δ0 trp1Δ63 ura3Δ0 met15Δ0 ho:: pTDH3-Cas9-tCYC1-TRP1 can1::PED ReSCuES[URA3-leu2]* | SparLox83. ReSCuES sequence inserted into JDY525 at an intergenic region in chr6. The *URA3* gene is in frame and the *LEU2* gene is not in frame. |
| JDY528 | a | *his3Δ200 leu2Δ0 lys2Δ0 trp1Δ63 ura3Δ0 met15Δ0 ho:: pTDH3-Cas9-tCYC1-TRP1 can1::PED ReSCuES[LEU2-ura3]* | Strain picked from nocodazole containing medium after SCRaMbLE induction in JDY526. The ReSCuES sequence is inverted so that the *LEU2* gene is in frame and the *URA3* gene is not in frame. |
| JDY529 | a | *his3Δ200 leu2Δ0 lys2Δ0 trp1Δ63 ura3Δ0 met15Δ0 ho:: pTDH3-Cas9-tCYC1-TRP1 can1::PED ReSCuES[LEU2-ura3]* | Strain picked from nocodazole containing medium after SCRaMbLE induction in JDY526. The ReSCuES sequence is inverted so that the *LEU2* gene is in frame and the *URA3* gene is not in frame. |
| JDY530 | a | *his3Δ200 leu2Δ0 lys2Δ0 trp1Δ63 ura3Δ0 met15Δ0 ho:: pTDH3-Cas9-tCYC1-TRP1 can1::PED ReSCuES[LEU2-ura3]* | Strain picked from rapamycin containing medium after SCRaMbLE induction in JDY526. The ReSCuES sequence is |

| | | | |
|---|---|---|---|
| | | | inverted so that the *LEU2* gene is in frame and the *URA3* gene is not in frame. |
| JDY531 | a | *his3Δ200 leu2Δ0 lys2Δ0 trp1Δ63 ura3Δ0 met15Δ0 ho:: pTDH3-Cas9-tCYC1-TRP1 can1::PED ReSCuES[LEU2-ura3]* | Strain picked from rapamycin containing medium after SCRaMbLE induction in JDY526. The ReSCuES sequence is inverted so that the *LEU2* gene is in frame and the *URA3* gene is not in frame. |
| JDY532 | a | *his3Δ200 leu2Δ0 lys2Δ0 trp1Δ63 ura3Δ0 met15Δ0 ho:: pTDH3-Cas9-tCYC1-TRP1 can1::PED ReSCuES[LEU2-ura3]* | Strain picked from benomyl containing medium after SCRaMbLE induction in JDY526. The ReSCuES sequence is inverted so that the *LEU2* gene is in frame and the *URA3* gene is not in frame. |
| JDY536 | a/alpha | *his3Δ200/his3Δ1 leu2Δ0/leu2Δ0 lys2Δ0/lys2Δ0 trp1Δ63/TRP1 ura3Δ0/ura3Δ0 met15Δ0/MAT15 ho:: pTDH3-Cas9-tCYC1-TRP1 can1::PED ReSCuES[URA3-leu2]* | JDY526 crossed with BY4742 |
| JDY541 | alpha | *his3Δ1 leu2Δ0 lys2Δ0 ura3Δ0 synIII HO::synSUP61* | Strain with complete *synIII* |
| JDY543 | a | *leu2Δ0 met15Δ0 ura3Δ0 his3Δ1* | ReSCuES inserted into BY4741. The *URA3* gene is in frame and the *LEU2* gene is not in frame. |
| JDY544 | a/alpha | *his3Δ200/his3Δ1 leu2Δ0/leu2Δ0 lys2Δ0/lys2Δ0 trp1Δ63/TRP1 ura3Δ0/ura3Δ0 met15Δ0/MAT15 ho:: pTDH3-Cas9-tCYC1-TRP1/ HO::synSUP61 can1::PED/CAN1 ReSCuES[URA3-leu2]* | JDY526 crossed with synIII |
| JDY546 | a/alpha | *leu2Δ0/leu2Δ0 met15Δ0/MAT15 ura3Δ0/ura3Δ0 his3Δ1/ his3Δ1 HO::synSUP61* | JDY543 crossed with synIII |
| JDY549 | a/alpha | *his3Δ200/his3Δ1 leu2Δ0/leu2Δ0 lys2Δ0/lys2Δ0 trp1Δ63/TRP1 ura3Δ0/ura3Δ0 met15Δ0/MAT15 ho:: pTDH3-Cas9-tCYC1-TRP1/ HO::synSUP61 can1::PED/CAN1 ReSCuES[LEU2-ura3]* | Strain selected after five SCRaMbLE rounds with JDY544. |

| | | | |
|---|---|---|---|
| JDY550 | a/alpha | *his3Δ200/his3Δ1 leu2Δ0/leu2Δ0 lys2Δ0/lys2Δ0 trp1Δ63/TRP1 ura3Δ0/ura3Δ0 met15Δ0/MAT15 ho:: pTDH3-Cas9-tCYC1-TRP1/ HO::synSUP61 can1::PED/CAN1 ReSCuES[LEU2-ura3]* | Strain selected after five SCRaMbLE rounds with JDY544. |
| JDY551 | a/alpha | *his3Δ200/his3Δ1 leu2Δ0/leu2Δ0 lys2Δ0/lys2Δ0 trp1Δ63/TRP1 ura3Δ0/ura3Δ0 met15Δ0/MAT15 ho:: pTDH3-Cas9-tCYC1-TRP1/ HO::synSUP61 can1::PED/CAN1 ReSCuES[LEU2-ura3]* | Strain selected after five SCRaMbLE rounds with JDY544. |
| JDY552 | a/alpha | *his3Δ200/his3Δ1 leu2Δ0/leu2Δ0 lys2Δ0/lys2Δ0 trp1Δ63/TRP1 ura3Δ0/ura3Δ0 met15Δ0/MAT15 ho:: pTDH3-Cas9-tCYC1-TRP1/ HO::synSUP61 can1::PED/CAN1 ReSCuES[LEU2-ura3]* | Strain selected after five SCRaMbLE rounds with JDY544. |
| JDY553 | a/alpha | *his3Δ200/his3Δ1 leu2Δ0/leu2Δ0 lys2Δ0/lys2Δ0 trp1Δ63/TRP1 ura3Δ0/ura3Δ0 met15Δ0/MAT15 ho:: pTDH3-Cas9-tCYC1-TRP1/ HO::synSUP61 can1::PED/CAN1 ReSCuES[LEU2-ura3]* | Strain selected after five SCRaMbLE rounds with JDY544. |
| JDY596 | a/alpha | *his3Δ200/his3Δ1 leu2Δ0/leu2Δ0 lys2Δ0/lys2Δ0 trp1Δ63/TRP1 ura3Δ0/ura3Δ0 met15Δ0/MAT15 ho:: pTDH3-Cas9-tCYC1-TRP1/ HO::synSUP61 can1::PED/CAN1 ReSCuES[LEU2-ura3]* | Strain selected after five SCRaMbLE rounds with JDY544. |
| JDY597 | a/alpha | *his3Δ200/his3Δ1 leu2Δ0/leu2Δ0 lys2Δ0/lys2Δ0 trp1Δ63/TRP1 ura3Δ0/ura3Δ0 met15Δ0/MAT15 ho:: pTDH3-Cas9-tCYC1-TRP1/ HO::synSUP61 can1::PED/CAN1 ReSCuES[LEU2-ura3]* | Strain selected after five SCRaMbLE rounds with JDY544. |
| JDY598 | a/alpha | *his3Δ200/his3Δ1 leu2Δ0/leu2Δ0 lys2Δ0/lys2Δ0 trp1Δ63/TRP1 ura3Δ0/ura3Δ0 met15Δ0/MAT15 ho:: pTDH3-Cas9-tCYC1-TRP1/ HO::synSUP61 can1::PED/CAN1 ReSCuES[LEU2-ura3]* | Strain selected after five SCRaMbLE rounds with JDY544. |
| JDY599 | a/alpha | *his3Δ200/his3Δ1 leu2Δ0/leu2Δ0 lys2Δ0/lys2Δ0 trp1Δ63/TRP1 ura3Δ0/ura3Δ0 met15Δ0/MAT15 ho:: pTDH3-Cas9-tCYC1-TRP1/ HO::synSUP61 can1::PED/CAN1 ReSCuES[LEU2-ura3]* | Strain selected after five SCRaMbLE rounds with JDY544. |

**Table S3 Sequences of edited sites**

| Site ID | Chromosome | Target sequence | Left homologous arm (HL) | Right homologous arm (HR) |
|---|---|---|---|---|
| I-1 | 1 | ATAAATGGCAC GTGTATGTA | AAAAAACAAAAGCAAATCACATGTGCACATACG TCCAGAATGATATCAAG | CTGTGTAAATATGATAATCATCTCGGACGAACGG CGTAGCACTCTCCATC |
| I-2 | 1 | GCGACAAACAT ACATATTAT | TACAGTCGTGATACGATTTACTGTCACTTAGCAAT AATATCTCGTACATA | ACACGTCGCCCTGAAAAAAAAAAACATAAGAAA AGAATACGAAAAAAAAG |
| I-3 | 1 | CCATTTCCAGT GCCTCCGAT | TTTGGCTTCCAGTATGCTTTCACGGAATTATTTCT CATGTACATTTAGCT | GAGGCATCATGGTACTACCGTGACGGAGAATACG TAGGCTGACTTTTTCG |
| II-1 | 2 | GAACGAACAG CAAGATAAAG | ACTTATCCATTATTTCCATCGTCAAAAAAAGGAA ATAAATACTGTTGCTC | AAACGAAAAGTAAGCAGCTTCCTCAATATGCCGT CAAACGTACGTTCGGG |
| II-2 | 2 | ACGACATGGAA GCGATACAC | AGAGTTTGGTTGCCACATGTACGCTTAGGACACT CATGACTATCACTCCT | ATACCCACTGGCCGATTCTCACCCTCTGCATTGTA TGATATTACTAGTAT |
| II-3* | 2 | N/A | N/A | N/A |
| II-4 | 2 | GACGGTAATGA TTAGAGTTT | AAAAATTTAACGTTCGATAATTAAGTAACAACAG AGAAAATATGAGCTTA | TGTTTTATTTACTGTCACCTTGATGAGCGACTAAA AAGATAGAACGCGGG |
| II-5 | 2 | GCGAAAGTGA AAGGTGCAAG | TACATAAAATTATATATAAGAAACACTTTTGCTTT AGCCTTCCTTTCTTT | CGTCCTTTTTCACTCACAGCAACAAGCAGCAAG CACTAAGTACGCAGTCA |
| II-6 | 2 | TCATAATGGCG ATGACAGGA | TGCTCGGACATGTTTGGGTGGACCTTATTCTACAT GTTTTACTTTTCTCG | GCAAGCTCATCTCCCCAGCTTTAAAGGGTAGCTC AAGGAACACCTACTTC |
| II-7 | 2 | CACTGCGCTGA AATATTACG | CACTTACGCTGCAATAAGCAGCAACGGCATTGTA GTGATCATTTTCTACG | GAAAAAAATTGAAAATTTTTACTCTTCTCGAGTG TTGAATCACTGCTGCG |
| II-8* | 2 | N/A | N/A | N/A |
| II-9 | 2 | CTCAGTTTCAA CATTATGAT | CTACTGCACTGTCATTATAGCCTAGTAAAGTATAT AGTGAATACAATATA | TAACTCCATCAGAAAATATATTCATCGTCATATAC GGAACATTCAGTTAT |

| | | | | |
|---|---|---|---|---|
| II-10 | 2 | CTAAAGATAAAACTAACTGC | GAAGACATTAATACCTTTATTCATATAAGCACTTTCATTATCATTTTTTA | CACCAAGAAGTTCGAAGCCAAGAGTAAAGATGTCAGGAACCGGATTATCG |
| II-11 | 2 | CCCATATAGTGATGCCTAAG | GCCTTGCTCGTCATGAGAACGACTAACAAGTAAGAGCGCGATGTTGCTGT | GCCAGGAATGGCAAATTTACTTGATAAACTTCAGGCGATTGGATTTTGGT |
| III-1 | 3 | GTTGTTTGAAGCCCTTTAAA | GTAGCAAAGTTAATCTGCCAATTGACAGTAGTTTAATATATGGTATTATC | AAAAAACGGGTTAGGGCCACCCGGCGCGAAGTAATAGCTGCTGATTGGTC |
| III-2 | 3 | TCGAGAGTTTCACCAACCAT | TTGAACAAACGGCTGAGACGGGCAATACATATGCTCTACTTCTTTTCCAT | GCATACATTACCTTACGTGTGTTAGTGTACTATATTATATATATATATAT |
| III-3 | 3 | GGATTGATTATATAGGCATA | CATTTATCTTCATATTCATGAATTTCCTTACTGGACCCCCACCTTAGCAT | GCATTCCGTCCACTGTATCGTAGGATTATTTTCCAACATTAGTTAACTTA |
| IV-1 | 4 | ATATTCTAGCTTCGTTGTCA | TTCCTAATTGTGCATTTTTTCAATAACAATACTTATTCATCCTTATAATT | GAACATAGCCCATACACCGCAGTTATTTATGATCATTTCGAACGGGAAGT |
| IV-2 | 4 | GTTTAACGGTGCAGTGAGTT | TAGATAATCTTACAAGGGACAAGTAGTCAAGCCTTGCTTATTATAATTCT | ACATTCATGCAACGTGGTAAATATGTGTCTCTTTGCTTCTGTATTTAAGC |
| IV-3 | 4 | AATAAGTCAGCCCCTCCCTT | ATTTTTTTTTTTCATTTTTTAAAGGGTTTCTCTACAGCCTACAGGCCTCC | AGTGCGCTGTTGACCTGCGTATATAAGAGGTATATCAGTGCCAGTAGGTA |
| IV-4 | 4 | CGGTGAGAGGTGAGAGGTGA | AGTATAACAGTATATCTGACACGCACGTGATGACCACGTAATCGCATCGC | CTGACTCAGCTTCACTAAAAAGGAAAATATATACTCTTTCCCAGGCAAGG |
| IV-5 | 4 | GATAGAGCAGTACTTATATA | CGAAGCCCCTTATCCCCTAGTTACCGAAGAAGGCCACCAATCTTAAGTTT | CTATATATAGACTGGTTCACAAGGTTATCAATATGAAACTTGCGCGATCA |
| IV-6 | 4 | CTATATCATAGCCAGTTAGC | AAAAAGCTACGCAAATATCGTATATCTGTTATACTACAAAACAATTACTT | AACGACTTCAGCTAAATGGACTATCCATGCTTTAGGCAGAGGCGAAGCGC |
| IV-7 | 4 | CAGAAGTAGATAAAGCAGCC | AAAAAAACAGGTAGAAGAAGGCTTGCTATAATTTGAACACTCTCTACCCT | TTGGCTTGAGAAACGTCATATCTATATATAGCGTAGATATGTTTATTCGC |

| | | | | |
|---|---|---|---|---|
| IV-8* | 4 | N/A | N/A | N/A |
| IV-9 | 4 | GGCCTTGATGT TATTAGTAC | TGGAAAAATATGACATAAGGTATGCGTATTAGTA AACTATTAGGGAGCCG | ATCTCACGCCGAGACTTACTTGGACTTTTCTCTAT TGTAAAGCGGAAAAA |
| IV-10 | 4 | CATCCTTACTAC TTTCCTCG | TTTTCGTTTCGCAGCGAATCCCTTTTAGCAGAGG AAAAAAAAGATGAAAC | AGACTCAACAGTAAAGGTTACTTTCAATTCAATA AACAAAAGGCACAGCG |
| IV-11 | 4 | AAAGGGAGCAT GTACATCTG | GGGAAAAGAATACTGCTACTGCTGTGCGAGACT TTGGTAGTAGGGATCGG | AATATATAAGCAGGAGCTCTCTACCTGGACCAAA TTGCCTTCTTATGTTG |
| IV-12, 13 | 4 | TCTCAGCATAG CATTAACAT | CGTAAGAAATACACATATAGTAGGTTTTGTCGCCT CCTCTTCCCCCTTTG | TGCCTCAGAGTGACAAAGAGAGAAATAGTTAAC TAGAATACGGTGCAAAT |
| V-1 | 5 | AGCGACTAACT ACCCTATTA | CTAATTATGTCGGCACATTTGAGAACCAATGGTT AGTTTCCAGCGCACCT | AGGGACGAAGGTGGTCTTTCTGAGGGAAGGAGG AAAAAAAGGTAAGAACC |
| V-2* | 5 | N/A | N/A | N/A |
| V-3 | 5 | CAAAGAGAGC ATGTCCATAG | CTACTCATTCAAAAATTATCCCTCTTCACTTCCCG TATTCACACTTGTTG | AAAAATGTAGTCTCACCCACAGAAAAGAAAAGA TCATTTGAAAATAAGAT |
| V-4 | 5 | ACCCATGTGAG ACTGAAACA | GAAAACCTTTGGAGAAAATCATTGCAAATTTAAA AGCTGTGCTTCAAAAA | TCCAGTGGTCTTCATCCGGACCGGTTCAAAGTCC TGCTCTACCTTCAATA |
| V-5 | 5 | TCTTAGATGCA GATATTCTT | TATAATTAGTTTTTCATCTGAAAGATATTTAGGGC ACCATTTTCTTTTGT | CGATTTACCCTGGTGGTACAGAAGATTATGTTAC ATAATTCATCAATTTT |
| V-6 | 5 | GGCAAATAGCT TCCTCTTTG | TTTGCGGAAGCTACTTTATTCCGGCCTGGAGTCA AAAGAGGAAGCTCGGT | CCGGGGCGCGGGGGGACGAGGCAAAAAGCAAA GAAAAGCAAAAAAAATAA |
| VI-1 | 6 | CCTTGCTGAAC ATTGAATAG | TAGAGGTTTATATTATAAAAGTGGAAAGGTAAAA TCAACAGCGTTTATTT | ACGGTAGAGACTACTATTGCTAAACAATTACTTA CATGGAAATGTACTGT |
| VI-2 | 6 | GTAATCACTATA AACGCGTA | AAGCATTGCTTCATGGAGGGGGTTGACTTCTTGA ATAAAATGGCTTTCTG | CGTGTTGCGTGGCTCTGATGATGGGCATTTCTAAT TTTAAGATCAACAAC |

| | | | | |
|---|---|---|---|---|
| VII-1 | 7 | ACGTCAAGTGAGAAGAGTTT | TTTAGTGAAGAAAGAGAAAAGTTGCTGCTCTAGATTTTGTATCGGCTATT | CCTTCGCTTCAGTTAGATCTTCTATTATTTCCTTTTTTTTCTTTTTGTTT |
| VII-2 | 7 | GAAGACCTATCAATTTTATG | TTTCTTTCTTTAGACATCAAACTGGTAGTTCTTATCAGTCTCAGCCTTTT | AGGGTAATGGTTGCTTCCTTTTCCCTAGCAGCGGCCCATCGCTTTAGCTT |
| VII-3 | 7 | ACTTATTAGTAAGAAGAGCA | TTCCGACTAAAACCCGCCTTCCCACGCGAAATTCTGGGCCGTTCAAGGCA | GGAAGGTACATTAAAGCAAGAGAACCGTGCATGAATTTATAGACATTTTT |
| VII-4 | 7 | AGCAAAAGTAGATCATTCAC | ATAAAACATAAACAAAAAAGAAAAATTAAGATTTGCAATTCTGCCGCTTA | AATATATATATATAAACGCATTTATAATCTTGTAACGTGCACTCAATTTA |
| VIII-1 | 8 | CACATCTTGCCTGTTATCTA | ACCTATGGGAAAGGTGTAACTCATCATTGCGCTTTCTACGGTACGGTTAT | CTCTATTCATCGGTGTTGCATGAAGATACGTCTTGTTCGCCAAGTATATA |
| VIII-2 | 8 | ACCTTAGCATTGGGATCAGC | GAGAGAATATCACTGTTTACGGCCCACAAAGAGAATGAGAAAATTGATAA | AGCCTTAGAATTCTGGGTGTGAACAGAAACACATTAGTGCATTAGTGTAT |
| VIII-3 | 8 | GAGCAGCGAGAACACGACCA | CTGCACTTTGCATCGGAAGGCGTTATCGGTTTTGGGTTTAGTGCCTAAAC | GCTATATAAATGGAAAGTTAGGACAGGGGCAAAGAATAAGAGCACAGAAG |
| IX-1 | 9 | GGCGCTATCAAAGGGAAACG | AACTTGACGCGTCAACATGAGGAGGGTAATGATGTGGTAGCGCCGTGTAA | ATAATAGTATTAACACCGCAGCTTTTTTTTCCTTTCTCCCTCTATTGGTT |
| IX-2 | 9 | CCAGGCAGAGTTGTGAAACC | AAATCAAAATAAACATCAAAAGAACCGCCAATTGATAAAAGCACAAAGTA | CCTCTTCTATTGTACCACACACAAATTTTTCTTTATTTAGGTAGAGGATG |
| X-1, 2 | 10 | TAGTAAAAGAAACGTCGATG | AATTATACCAACATGGTTGTAGCATTTCAAGTTAGCTTGTTCGAGCTGTA | CTAATTTGTTCCGGTGACTTTCAGGTCACGTGTTTTCCATGCGTGCGTTG |
| X-3, 4 | 10 | ACTCTCAACAGTGATATTTC | AACAACTCATTACTAGACTACTACCAGTTACTACACATCGATACAACTCC | TCATGTCAAGACGTAGTATAGTATTGCAACAGGAAAAAAAAATCTTATTG |
| X-5 | 10 | CAACGGACCCTCTTAATTAG | GGTTTTTTTCGCCTCTCTTCAAGTTTCCTATACCCGAGCTTAACACAACA | TATCCTCCGTTCTTTTCTTCCGCTTTCCTAATCGAGATACCAGAGCACGG |

| | | | | |
|---|---|---|---|---|
| X-6 | 10 | GATATGTCTTGCAAAACTAT | CCTCCCCGTTTGTATCAAAAGTATCCGAAACTGTTCTATGCGTGCAAAAT | AAAGACCCTTCCACTACAGGTACAGTTTAGAAATGCCGGAGCTGCAAGTA |
| X-7 | 10 | CACCACTGTCTTCTTTCCTG | AATCCCCTAAACATTCAGATTGTAAACTAGGGTTGAGAAAATGACTCATC | CATTCTATAGATTATTGTGAATGACTCTTATTGATGAGATGGCAATAACT |
| XI-1 | 11 | CATGGATATTAGAGGATAGT | CATTATTTATATCATATTCTTACACTTCATACAATTATATATACTCGTAC | TTTTTGTTTAAACTAATTCTATGTAACCTCGATGGCATGAGTCCATAATA |
| XI-2 | 11 | ACGATGCGATGATAAATATT | ACATGCAAAATCAGCCCACTCGAAGTTAGATACTGCGTAGACGGATTTGC | GACAGTGCGTGCCCACCCATAAGTTGCTGTAGAGCTTACAAATTGTAAAG |
| XI-3 | 11 | GAAAAGCCTCAACTATTTAT | TGTAGTGACGCGGAAACGTTTTACTCGTCAATGGACATTTGCGAATGTGT | CTGCATTCTGTCAAGAAGGGTATGTGTATGAACATGCAAATGACACTGTA |
| XI-4 | 11 | TTGTGCGCTGAGTAATCATT | AAAAAAGGGATTGACATTTCTTCGAGAATTAGTTGAGAAACCCTTTTAGA | TTGATTATAAGCTCTGAAAGGTTACTGCTACTGCTAGTAGTCTTTCAGGA |
| XII-1 | 12 | GAATGAGGGATGGAACAAGA | CCCCATTTTCCCGAATTTTCTTCTACGTTTTCTTTTTTCAGAAATACTTC | AAGCGACATACAAAAACTGAGTCAGTTTAGTCTTAATAGGTGCAATATTT |
| XII-2 | 12 | GAAGTGGTGTAGTAGTGATG | TATAGTATATACTACCTGTAAATATGTGCGATGCACAATTAACATTACCT | TACTGCTAGCACTGTCCTCTTGTGCTTGGCCCCTTAAGAGTGTTCTAAGA |
| XII-3 | 12 | TACTGCACTGTCACTTACCA | AAGTTTTATAAGCATTTTTATGTAACGAAAAATAAATTGGTTCATATTAT | AAAGACCAGACAAGAAGTTGCCGACAGTCTGTTGAATTGGCCTGGTTAGG |
| XII-4 | 12 | TCTTCTTGACAGCACTGCCT | AATGCCATTCTGCCTAGCCCATTTGCGTCTTCGCTGCCGTAACCATTCTC | TCAGTTCTCTATATTTGTCGCGCCGCCGCAAAATGCACCCACATTAAAGA |
| XII-5 | 12 | GATGCATTACGAGAAGGTTA | TTGTTCGTTGTGCACGTAGGATGTATATTGAACAAGCATGACCAGAATCT | GATGATATCAGACCTCCGAAGTCCATGTTGCAAAATGTGCCGACTTTCCG |
| XII-6 | 12 | AATTTAAAACCGTGGCTTGC | GCCGCAAATACAGAGGCGCCCCAGACAACACCGCAGTGTGAAGCACTGTC | AGATGCCCAGACCAACCCTGTTGGGTTTTTCTCTCGAGCACGCCGTTATA |

| | | | | |
|---|---|---|---|---|
| XII-7 | 12 | TTGCATACAAATCCCTCTGA | AGTGGAAAACGACATAAATAATAATTTTATAATATAATAATGATAATTCA | ATGATTGATGACGCGAGAAAAAAAAACGCGAAATTTTTCTTCCCAAAGCT |
| XII-8 | 12 | CTACTTGTCGATGCATATAT | ATAAAAAAGCAATATTTTTTTGCGAGCTATTTAGTGATATAGCCGCCCAG | CTGAGAGTACTAATACAATTATCGGAAGACCTCAAAGTAAAATATAGAAG |
| XIII-1, 2, 3, 4 | 13 | CGTTAAACGCTGCTGATTGA | CATTAGAAATAAGGCTCTCGTTGATCATCCTTGTAACTGAAAATTAGAAC | ATTCCAGCCGTTCTCTTCCAACCCCTTTTACCCCGATTGTTCGTCCACTA |
| XIII-5 | 13 | CAACCATCTTCGCCAAGTAG | ATCATACTTAGTAACAAGAAAGACAAAAGCGCAAACCGAACCGCCCAGCT | CAAACTTATATAAGCTCCAACGATGTCCCCATCAATTAAGAACCCTCGAT |
| XIII-6 | 13 | GTTTAGCTGTTATCCTCTGA | GTTCGGCAGTCTAAAGGTTAAACATTATCAGAAATTATCTATTCCTATTT | TTGAATTGTAATTTTACTAAGAAAAGGAGAGGAAAAGGACGTGCATATAT |
| XIII-7 | 13 | CTTTATCCATCATTGAGACA | CGTGCTTTTATTAGTGGGCCCCTTCTTTGAGACCCCGCGGGTGATATGGC | TTGTTGGAAGCAATTACCAAGACAATCACAAACGAAGGTTGCTACCGAAG |
| XIV-1 | 14 | AATAGTTTGAGCCAGCACGA | CGATGATAAATCCTCCGCCGCATGATGCTTTTGATTTGCCTAAGGGCCTG | GAGGGTCAACATACCTTGAAAATCCAAGTAAAAGGATGGATATCGTTATA |
| XIV-2 | 14 | ATATATAGCTCGCTCATGCA | TGAGAGGGTAGGCAAGAAGGTCGAAAGGAAAACAAAAAAACGTATCGCTG | CACAAATACAGGGGTAGGGTCCTCGACGTAGTAGACATTCTGCTTCTATA |
| XIV-3 | 14 | CTGTGACGTCGATGCATGCG | GACACTTATACTTGGTGGGGAATCGCCCGTCAGGCCTGAACGCAACGAAC | TGAGCTCAGGCCGCATCACGGCCGTTACGCCCTCCAGAGTCACCACGACT |
| XV-1 | 15 | CGATATGATGGTGATGGTGA | AGCAATTCGGGAGGGCGAAAAATAAAAACTGGAGCAAGGAATTACCATCA | CCTTAGCCTCTAGCCATAGCCATCATGCAAGCGTGTATCTTCTAAGATTC |
| XV-2, 3 | 15 | CTGTGAGCAGTAATTATCAA | AACCATGAAAATATGGTCTCCAGGTTATCAATAGCTTCGAATTGGAAATA | ATGCTATTATATAAATATACATACCTACACCCATCCCATATTTACATAGA |
| XV-4 | 15 | GCTCTATCGGTCCGCTAGCT | GTAAGACCGATCCACTTTGCCAGCTGCTTACGCTGCGGAAAGTAAACAGA | CGGCTGAACTTTATAACAAATGCGCCTTCTAACAAGCGATGAAGCCATGC |

| | | | | |
|---|---|---|---|---|
| XVI-1 | 16 | GTGGACGGTTCTTAAAATTC | GGATTGTTTACCAGCCGGCAGGACTCTGTTGATTTGTTTCACCTGTGATG | CACCTGTTAATAATATAATTGTGCAAATGCGCGCTTTTTTCGCCGCTCAC |
| XVI-2 | 16 | ATAACCACCATGTCAGCACC | TCCTGCTGGTGTCGTATTATTTTTTGAAATTATTTTTCAATAACCACCAT | TTCTTTTACAATTATACAAACACACATATCTCAAAATCACTCAAGAGGTC |

*Off-target site.

**Table S4 Primer sequences.**

| Systematic name | Other name | Sequence | Description |
|---|---|---|---|
| MGO001 | SHO582 | CACCTTACTGGAATTTAGTCCCTGCTA | Used to verify loxPsym insertion, paired with site-specific primers. |
| MGO002 | XIII-5_R | CTTCGCCAACTGCAACGGAATA | Used to verify duplication. For amplicon 1 in Extended Data Fig 3a. |
| MGO003 | XIII-6_F | GGCGAATTTGACACAGCTAGTAAGG | Used to verify duplication. For amplicon 1 in Extended Data Fig 3a. |
| MGO004 | IV-1_R | GGCAGCATCACCAATCAATCTTTCT | Used to verify duplication. For amplicon 2 in Extended Data Fig 3a. |
| MGO005 | IV-2_F | GCGGAAACCAGCGTCACTAATTT | Used to verify duplication. For amplicon 2 in Extended Data Fig 3a. |
| MGO006 | IV-9_R | CTTAATGGAGATTGAGGCAGCAAT | Used to verify translocation. For amplicon 3 Extended Data Fig 3a. |
| MGO007 | XII-6_F | GTCATCCTCCCTGTGTTTACAGTG | Used to verify translocation. For amplicon 3 Extended Data Fig 3a. |
| MGO008 | X-4_R | TTCTATAGAACGTGTATGGCTCACG | Used to verify translocation. For amplicon 4 Extended Data Fig 3a. |
| MGO009 | VII-2_F | GCAGGTGTTAATAGAGATGAAGGTAGATCC | Used to verify translocation. For amplicon 4 Extended Data Fig 3a. |
| MGO010 | X-5_R | CTCTTCACCAATCAGGGACATATCAGTACT | Used to verify inversion. For amplicon 5 in Extended Data Fig 3a. |

| | | | |
|---|---|---|---|
| MGO011 | X-6_F | CGCTATCTCAAGGCAAACGTTCT | Used to verify inversion. For amplicon 5 in Extended Data Fig 3a. |
| MGO012 | I-3_R | CCGAGCTAATTTCAAGTGGGTGAC | Used to verify inversion. For amplicon 6 in Extended Data Fig 3a. |
| MGO013 | I-2_F | CTGTACATACCGAAACGCCATCA | Used to verify inversion. For amplicon 6 in Extended Data Fig 3a. |
| MGO014 | IV-5_F | TTCCTTCTTGTCGACGACAGGTTTC | Primer A in Extended Data Fig 6b. Used to verify translocation in JDY528 strain. |
| MGO015 | IV-5_R | CAGGCGTTCTTGTATCTAGTAATCTCCT | Primer B in Extended Data Fig 6b. Used to verify translocation in JDY528 strain. |
| MGO016 | XIV-1_R | ATTTGGAAAATTTACCACTGCCCATGG | Primer C in Extended Data Fig 6b. Used to verify translocation in JDY528 strain. |
| MGO017 | IV-9_R | CTTAATGGAGATTGAGGCAGCAAT | Primer D in Extended Data Fig 6b Used to verify duplication in JDY528 strain. |
| MGO018 | IV-7_F | GCACATTGAATTTACACTCCCGATC | Primer E in Extended Data Fig 6b. Used to verify duplication in JDY528 strain. |
| MGO019 | LCO1268 | ACCCTTACCATCGCAGGACTTTTCAGTC | Primer R1 used in Extended Data Fig 7c. |
| MGO020 | LCO1269 | CGGTACTACCAGCGTTTTGTTGTTGGTC | Primer F1 used in Extended Data Fig 7c. |
| MGO021 | LCO1270 | TAGCGTCTTTCATCGAGGTAGCGTTTGC | Primer R2 used in Extended Data Fig 7c. |
| MGO022 | LCO1271 | GTTGCCGAAGAGACACCAAAATGTGCC | Primer F2 used in Extended Data Fig 7c. |
| MGO023 | LCO1272 | GATACAAATGCCCCGGAGAATCTAGTGTACC | Primer R3 used in Extended Data Fig 7c. |
| MGO024 | LCO1273 | AACAACTCGGCTGCCGTCTGGGCTATAA | Primer F3 used in Extended Data Fig 7c. |

**Table S5 Main software and algorithms used in this paper.**

| Software and algorithms | Source | Identifier |
|---|---|---|
| MinKNOW | Oxford Nanopore Technologies | MinION software (MinKNOW) upgraded to enable increased data yield, other benefits (nanoporetech.com) |
| NanoFilt | De Coster, W., D'Hert, S., Schultz, D.T., Cruts, M., and Van Broeckhoven, C. (2018). NanoPack: Visualizing and processing long-read sequencing data. Bioinformatics 34, 2666–2669. | https://github.com/wdecoster/nanofilt |
| sniffles | Sedlazeck, F.J., Rescheneder, P., Smolka, M., Fang, H., Nattestad, M., Von Haeseler, A., and Schatz, M.C. (2018). Accurate detection of complex structural variations using single-molecule sequencing. Nat. Methods 15, 461–468. | https://github.com/fritzsedlazeck/Sniffles |
| NGMLR | Sedlazeck, F.J., Rescheneder, P., Smolka, M., Fang, H., Nattestad, M., Von Haeseler, A., and Schatz, M.C. (2018). Accurate detection of complex structural variations using single-molecule sequencing. Nat. Methods 15, 461–468. | GitHub - philres/ngmlr: NGMLR is a long-read mapper designed to align PacBio or Oxford Nanopore (standard and ultra-long) to a reference genome with a focus on reads that span structural variations |
| Canu | Koren, S., Walenz, B.P., Berlin, K., Miller, J.R., Bergman, N.H., and Phillippy, A.M. (2017). Canu: Scalable and accurate long-read assembly via adaptive κ-mer weighting and repeat separation. Genome Res. 27, 722–736. | https://github.com/marbl/canu |
| BWA-mem | Li, H. (2013). Aligning sequence reads, clone sequences and assembly contigs with BWA-MEM. 00, 1–3. | https://github.com/lh3/bwa |
| GATK | | https://gatk.broadinstitute.org/hc/en-us |

| | | |
|---|---|---|
| Cutadapt | Kechin, A., Boyarskikh, U., Kel, A., and Filipenko, M. (2017). CutPrimers: A New Tool for Accurate Cutting of Primers from Reads of Targeted Next Generation Sequencing. J. Comput. Biol. 24, 1138–1143. | https://cutadapt.readthedocs.io/en/stable/ |
| HISAT2 | Kim, D., Paggi, J.M., Park, C., Bennett, C., and Salzberg, S.L. (2019). Graph-based genome alignment and genotyping with HISAT2 and HISAT-genotype. Nat. Biotechnol. 37, 907–915. | https://github.com/DaehwanKimLab/hisat2 |
| Picard | | https://github.com/broadinstitute/picard |
| Htseq-count | Anders, S., Pyl, P.T., and Huber, W. (2015). HTSeq-A Python framework to work with high-throughput sequencing data. Bioinformatics 31, 166–169. | https://htseqhedocs.io/en/master/ |
| DEseq2 package in R. | Love, M.I., Huber, W., and Anders, S. (2014). Moderated estimation of fold change and dispersion for RNA-seq data with DESeq2. Genome Biol. 15, 1–21. | https://bioconductor.org/packages/release/bioc /html/DESeq2.html |
| LAST | Kiełbasa, S.M., Wan, R., Sato, K., Horton, P., and Frith, M.C. (2011). Adaptive seeds tame genomic sequence comparison. Genome Res. 21, 487–493. | https://gitlab.com/mcfrith/last |
| Minimap2 | Li, H. (2018). Minimap2: Pairwise alignment for nucleotide sequences. Bioinformatics 34, 3094–3100. | https://github.com/lh3/minimap2 |
| Cytoscape | Shannon, P., Markiel, A., Ozier, O., Baliga, N.S., Wang, J.T., Ramage, D., Amin, N., Schwikowski, B., and Ideker, T. (2003). Cytoscape: A Software Environment for Integrated Models of Biomolecular Interaction Networks. Genome Res. 13, 2498–2504. | https://cytoscape.org/ |
| Stringtie | Kovaka, S., Zimin, A. V., Pertea, G.M., Razaghi, R., Salzberg, S.L., and Pertea, M. (2019). Transcriptome assembly from long-read RNA-seq alignments with StringTie2. Genome Biol. 20, 1–13. | https://github.com/gpertea/stringtie |
| Distiller-nf | | https://github.com/open2c/distiller-nf |

| | | |
|---|---|---|
| Juicer | Durand, N.C., Shamim, M.S., Machol, I., Rao, S.S.P., Huntley, M.H., Lander, E.S., and Aiden, E.L. (2016). Juicer Provides a One-Click System for Analyzing Loop-Resolution Hi-C Experiments. Cell Syst. 3, 95–98. | https://github.com/aidenlab/juicer |
| Pastis | Varoquaux, N., Ay, F., Noble, W.S., and Vert, J.P. (2014). A statistical approach for inferring the 3D structure of the genome. Bioinformatics 30, i26–i33. | https://github.com/hiclib/pastis |
| Pymol | L DeLano, W. (2002). Pymol: An open-source molecular graphics tool. {CCP4} Newsl. Protein Crystallogr. 40, 1–8. | https://github.com/schrodinger/pymol-open-source . |
| Fastp v0.12.4 | Chen, S., Zhou, Y., Chen, Y., and Gu, J. (2018). Fastp: An ultra-fast all-in-one FASTQ preprocessor. Bioinformatics 34, i884–i890. | https://github.com/OpenGene/fastp |
| Samtools v1.7 | Danecek, P., Bonfield, J.K., Liddle, J., Marshall, J., Ohan, V., Pollard, M.O., Whitwham, A., Keane, T., McCarthy, S.A., Davies, R.M., et al. (2021). Twelve years of SAMtools and BCFtools. Gigascience 10, 1–4. | https://github.com/samtools/samtools |
| Pilon | Walker, B.J., Abeel, T., Shea, T., Priest, M., Abouelliel, A., Sakthikumar, S., Cuomo, C.A., Zeng, Q., Wortman, J., Young, S.K., et al. (2014). Pilon: An integrated tool for comprehensive microbial variant detection and genome assembly improvement. PLoS One 9. | https://github.com/broadinstitute/pilon |

**Table S6 The numbers of rearrangement for strains in Figure 4.**

| Strain | Cre/loxPsym mediated rearrangement* | | | Homologous recombination mediated rearrangement |
|---|---|---|---|---|
| | SynIII derived | SparLox83 derived | SynIII and SparLox83 derived | |
| JDY549 | 25 | 1 | 0 | 0 |
| JDY550 | 8 | 1 | 1 | 1 |
| JDY551 | 15 | 1 | 0 | 0 |
| JDY552 | 19 | 1 | 0 | 0 |
| JDY553 | 16 | 0 | 0 | 0 |
| JDY596 | 23 | 1 | 0 | 0 |
| JDY597 | 15 | 1 | 0 | 0 |
| JDY598 | 11 | 0 | 0 | 0 |
| JDY599 | 15 | 1 | 0 | 0 |

***Rearrangements were counted by structural variatiant juctions.**
